# Epitope-based peptide vaccine design and target site characterization against novel coronavirus disease caused by SARS-CoV-2

**DOI:** 10.1101/2020.02.25.965434

**Authors:** Lin Li, Ting Sun, Yufei He, Wendong Li, Yubo Fan, Jing Zhang

## Abstract

The outbreak of the 2019 novel coronavirus (SARS-CoV-2) has infected thousands of people with a large number of deaths across 26 countries. The sudden appearance of the virus leads to the limited existing therapies for SARS-CoV-2. Therefore, vaccines and antiviral medicines are in desperate need. This study took immune-informatics approaches to identify B- and T-cell epitopes for surface glycoprotein (S) of SARS-CoV-2, followed by estimating their antigenicity and interactions with the human leukocyte antigen (HLA) alleles. We identified four B cell epitopes, two MHC class-I and nine MHC class-II binding T-cell epitopes, which showed highly antigenic features. Allergenicity, toxicity and physiochemical properties analysis confirmed the specificity and selectivity of epitopes. The stability and safety of epitopes were confirmed by digestion analysis. No mutations were observed in all the selected B- and T-cell epitopes across all isolates from different locations worldwide. Epitopes were thus identified and some of them can be potential candidates for vaccine development.

## Introduction

The SARS-CoV-2 (coronavirus disease 2019; previously 2019-nCoV) has recently emerged as a human pathogen leading to 51,000 confirmed cases globally and at least 1,600 deaths [1]. SARS-CoV-2 virus is an enveloped, positive single-stranded RNA coronavirus with the genome size of approximately 29.9 kb. SARS-CoV-2 is closely related to several bat coronaviruses and the SARS-CoV virus [2, 3], and all belong to the B lineage of the beta-coronaviruses [4]. The transmission of SARS-CoV-2 appears to contain the way from human to human and from contact with infected surfaces and objects, causing WHO declare a Public Health Emergency of International Concern (PHEIC) on January 30^th^, 2020 [5–7].

Structural proteins are important targets for vaccine and anti-viral drug development due to their indispensable function to fuse and enter into the host cell [8]. SARS-CoV-2 utilizes glycosylated spike (S) protein to gain entry into host cells. The S protein is a trimeric class I fusion protein and exists in a metastable prefusion conformation that undergoes a dramatic structural rearrangement to fuse the viral membrane with the host-cell membrane [1, 9, 10]. The S protein includes the receptor binding S1-subunit and the membrane fusion S2-subunit. The S1 subunit receptor-binding domain (RDB) is specifically recognized by the host receptor. When the S1 subunit binds to a host-cell receptor, the prefusion trimer is destabilized, resulting in the shedding of S1 subunit, and the state transition of S2 subunit to a stable postfusion conformation [11]. The critical function of the S protein can be a breakthrough in vaccine design and development.

Great efforts are being made for the discovery of antiviral drugs, but there are no licensed therapeutic or vaccine for the treatment of SARS-CoV-2 infection available in the market. Developing an effective treatment for SARS-CoV-2 is therefore a research priority. It is time-consuming and expensive to design novel vaccines against viruses by the use of kits and related antibodies [12]. Thus, we chose the method of immune-informatics, which is more efficient and more applicable for deep analysis of viral antigens, B- and T-cell linear epitope prediction, and evaluation of immunogenicity and virulence of pathogens. Among those can be analyzed, B-cell can recognize and activate defense responses against viral infection, T-cell and antibody reactions may recover extreme respiratory infection.

In this manuscript, we applied immuno-informatics approach to identify potential B- and T-cell epitopes based on the S protein of SARS-CoV-2. The antigenicity of all the epitopes were estimated and the interactions with the human leukocyte antigen (HLA) alleles were evaluated for MHC class-I epitopes. Allergenicity, toxicity, stability and physiochemical properties were also investigated for exploring the antigenicity, stability and safety of the identified epitopes. The conservation of all B- and T-cell epitopes were examined across all isolates from different locations. Some of these identified epitopes could be used as promising vaccine candidates.

## Methods

### Data retrieval and structural analysis

Primary sequence of SARS-CoV-2 protein was retrieved from NCBI database using accession number MN908947.3 [3]. Experimentally known 3D structure of SARS-CoV-2 S protein was retrieved from Protein Data Bank (PDB ID: 6VSB)[1]. Protein sequence was analyzed for its chemicals and physical properties including GRAVY (Grand average of hydropathicity), half-life, molecular weight, stability index and amino acid atomic composition via an online tool Protparam [13]. TMHMM v2.0 (http://www.cbs.dtu.dk/services/TMHMM/) was applied to examine the transmembrane topology of S protein. Secondary structure of SARS-CoV-2 protein was analyzed by PSIPRED [14]. Existence of disulphide-bonds were examined through an online tool DIANNA v1.1 which uses trained neural system to make predictions [15]. Antigenicity of full-length S protein was evaluated by vaxijen v2.0 [16].

### B-cell epitope prediction

IEDB (Immune-Epitope-Database And Analysis-Resource) [17] with default parameter settings were used to predict B-cell epitopes. Epitopes predicted by linear epitope prediction of Bepipred and Bepipred2.0, Kolaskar and Tongaonkar antigenicity, Parker hydrophilicity, Chou and Fasman beta turn, and Karplus and Schulz flexibility. BcePred [18] was also used to predict B-cell epitopes using accessibility, antigenic propensity, exposed surface, flexibility, hydrophilicity, polarity and turns. Predicted B-cell epitopes by IEDB and BcePred were combined to the B-cell epitope candidate list. Based on transmembrane topology of S protein predicted by TMHMM v2.0, only epitopes on the outer surface were remained, and other intracellular epitopes were eliminated. VaxiJen 2.0 [16] was applied to evaluate the antigenicity of the remained epitopes. A stringent criteria was used to have antigenicity score of 0.9 and 14 residue length epitopes viewed adequate to start a defensive immune reaction. A B-cell discontinuous epitope forms the antigen-binding interface through fragments scattered along the protein sequence. DiscoTope2.0[19] with discotope score threshold of −3.7 was used to predict discontinuous epitopes. Pymol was used to examine the positions of selected linear and discontinuous epitopes on the 3D structure of SARS-CoV-2 protein.

### T-cell epitope prediction

Cytotoxic T-lymphocyte epitopes are important in developing vaccine. Peptide_binding_to_MHC_class_I_molecules tool of IEDB and HLA class I set [20] was utilized to predict MHC class I binding T-cell epitopes for S protein. Peptide_binding_to_MHC_class_II_molecules tool of IEDB and HLA class II set [21] was utilized to predict T-cell epitopes for S protein. Percentile rank with threshold of 1% for MHC class I binding epitopes and 10% for MHC class II binding epitopes were used to filter out peptide-allele with weak binding affinity. The antigenicity score of each epitope was calculated by VaxiJen v2.0. A high stringent standard was used to filter peptides with antigenicity score larger than or equal to 1, the number of binding alleles larger than or equal to 3 for MHC class I binding epitopes and 5 for MHC class II binding epitopes.

### Characterization of selected B-cell and T-cell epitopes

All selected B-cell and MHC class I and II binding T cell epitopes were examined for their allergenicity, hydro and physiochemical features, toxicity and digestion. Allergenicity of B-cell and T-cell epitopes were assessed by Allergen FP 1.0 (http://ddg-pharmfac.net/AllergenFP/). Toxicity of B-cell and T-cell epitopes along with hydrophobicity, hydropathicity, hydrophilicity and charge were evaluated by ToxinPred (https://webs.iiitd.edu.in/raghava/toxinpred/index.html). The peptides that can be digested by several enzymes are usually non-stable, while the peptides digested by fewer enzymes are more stable, so that those are more favorable vaccine candidates. Examined by protein digest server (http://db.systemsbiology.net:8080/proteomicsToolkit/proteinDigest.html), the digestion of B- and T-cell epitopes by 13 enzymes including Trypsin, Chymotrpsin, Clostripain, Cyanogen Bromide, IodosoBenzoate, Proline Endopept, Staph Protease, Trypsin K, Trypsin R, AspN, Chymotrypsin (modified), Elastase, and Elastase/Trypsin/Chymotryp.

### Protein-epitope interaction evaluation

The 3D structure of human HLA-B35:01(PDB ID: 1A9E) at a resolution of 2.5 Å, HLA-B*51:01 (PDB ID: 1E27) at a resolution of 2.2 Å and HLA-B*53:01 (PDB ID: 1A1O) at a resolution of 2.3 Å were downloaded from protein databank (RCSB PDB) and used for evaluating their interactions with selected epitopes. Protein-peptide interactions were performed by PepSite [22] with the top prediction chosen from a total of 10 epitope-protein interaction reports.

### Conservation analysis of selected B- and T-cell epitopes

The S protein sequences were taken from an open access database NGDC (https://bigd.big.ac.cn/ncov/), where 134 SARS-CoV-2 virus strain sequences are documented from 38 locations worldwide. The mutations in the S protein are documented in 37 isolates. The phylogenetic tree of the S proteins with mutations were generated using MEGA software [23]. By performing the multiple-sequence-alignment against the S protein sequences collected from different locations, all the selected epitopes were examined for their variability and conservation.

## Results

### Structural analysis of SARS-CoV-2 S protein

The physiochemical properties of SARS-CoV-2 S protein calculated by Protparam demonstrate that it contains 1273 amino acids (aa) with molecular weight of 141.18 kDa. Of the 1273 residues, 110 aa and 103 aa were found as negatively and positively charged, respectively. Theoretical iso-electric point (PI) of S protein was 6.24, reflecting its negative nature. The instability-index (II) was computed to be 33.01, which categories the S protein as stable. Aliphatic-index was 84.67 with GRAVY (grand average of hydropathicity) value of −0.079, which reveals a thought of proportional volume hold by aliphatic side chain. Half-life of the S protein was estimated as 30 hours for mammalian reticulocytes, > 20 hours for yeast, > 10 hours for Escherichia coli, which measures the total time taken for its vanishing after it has been synthesized in cell. The number of Carbon (C), Oxygen (O), Nitrogen (N), Hydrogen (H), and Sulfur (S) of a total of 19710 atoms were formulated as C_6336_H_9770_N_1656_O_1984_S_54_. The details of the physiochemical properties of SARS-CoV-2 S protein can be seen in Supplementary Table 1.

Secondary structure of S protein were generated by PSIPRED [14], showing that Beta strand (26.3%), Helixes (24.4%), and coil (49.3%) are present in structure (Supplementary Figure 1). 20 disulfide (S-S) bond positions were identified by DiANNA (Supplementary Table 2). 40 cysteine residues were identified by DiANNA in the full-length of the S protein sequence, which made 20 disulfide (S-S) bonds at the following positions (15-1240, 131-391, 136-662, 166-1236, 291-671, 301-336, 361-488, 379-743, 432-1235, 480-1248, 525-1247, 538-1043, 590-617, 649-1241, 738-1243, 749-1126, 760-1250, 840-1032, 851-1254, 1082-1253) (Supplementary Table 2). Antigenicity analysis of the full-length protein confirmed that S protein was an expected antigen with antigenicity score of 0.4646 by Vaxijen. The transmembrane protein topology predicted by TMHMM showed that residues from 1 to 1213 were exposed on the surface, residues from 1214 to 1236 were inside transmembrane-region and residues from 1237 to 1273 were within the core-region of the S protein (Fig. 1A).

**Figure 1.**
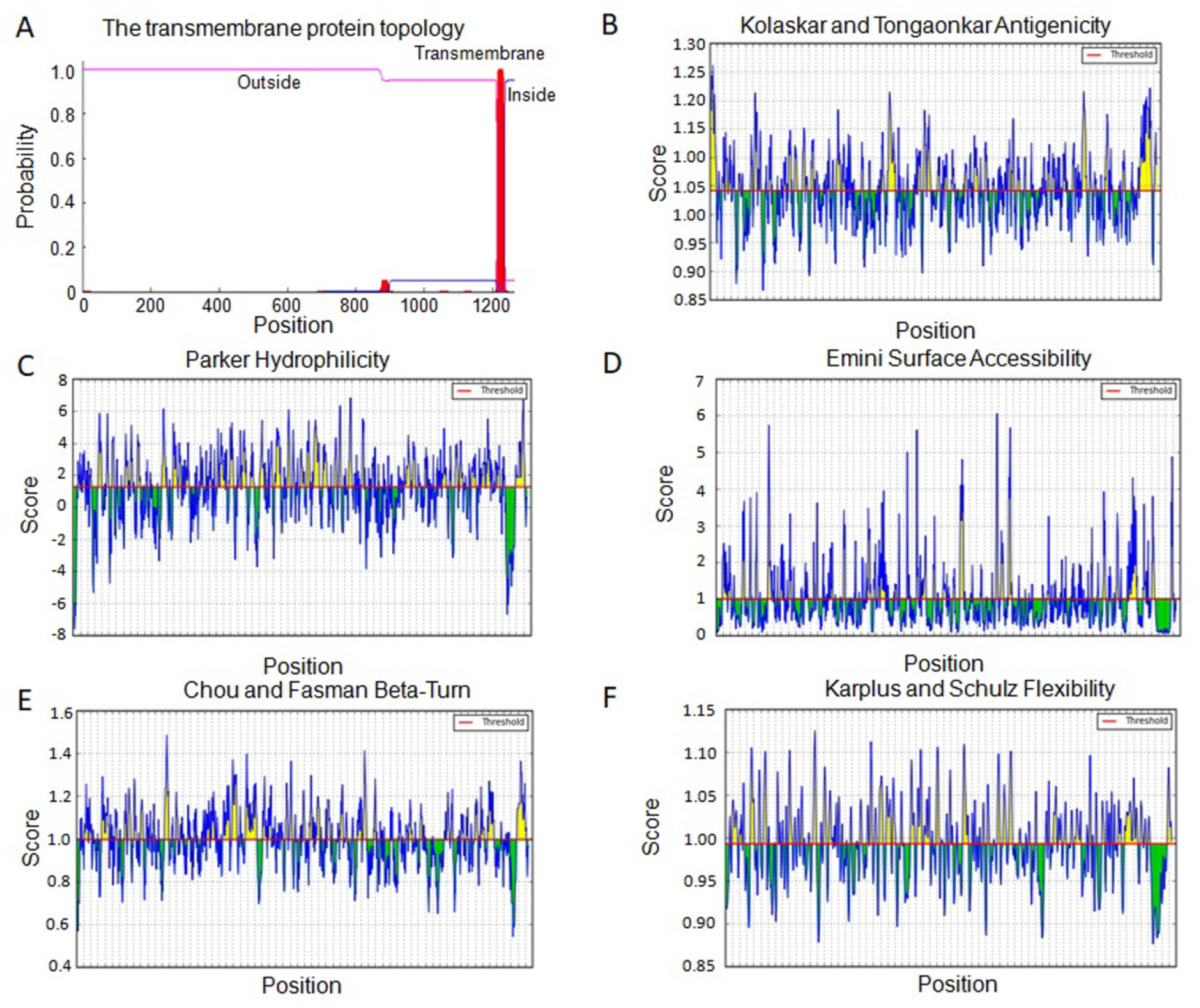
A.The transmembrane protein topology predicted by TMHMM; B.Antigenicity prediction using Kolaskar and Tongaonkar antigeicity scale; C.Hydrophilicity prediction using Parker hydrophilicity; D.Surface accessibility by Emini surface accessibility; E.Beta turns prediction by Chou and Fasman beta turn; F.Flexibility by Karplus and Schulz flexibility.

### Identification of B-cell epitopes

B-cell epitopes can guide B-cell to recognize and activate defense responses against viral infection. Recognition of B-cell epitopes depended on predictions of linear epitopes, antigenicity, hydrophilicity, accessibility of surface, beta-turn and flexibility [24]. B-cell epitopes of S protein were predicted using IEDB [17]. A total of 23 and 26 linear epitopes (Supplementary Table 3a) were identified by Bepipred and by Bepipred2.0, respectively. Kolaskar and Tongaonkar antigenicity of S protein was analyzed with default parameter settings by assessing the physiochemical properties of the amino acid and their abundance in known B-cell epitopes. The antigenic tendency value of S protein was estimated to be 1.041 (average), 0.866(minimum) and 1.261 (maximum) (Fig. 1B). Hydrophilic region is important in initiating immune response, which is generally uncovered on the surface of protein. Parker hydrophilicity of S protein was found to be 1.238 (average), −7.629 (minimum) and 7.743 (maximum) (Fig. 1C). To find the surface availability of B-cell epitopes, Emini surface accessibility was predicted for S protein with average of 1, minimum of 0.042, and maximum of 6.051 (Fig. 1D). Exposed on the surface and hydrophilic in nature making beta turn a vital structure in starting the defense response. Therefore, we predicted Chou and Fasman beta turn to gain the result, 0.997 (average), 0.541 (minimum) and 1.484 (maximum) in S protein (Fig. 1E). As the parts of epitope connecting with antibodies are typically elastic in nature, we predicted Karplus and Schulz flexibility of S protein and the result was 0.993 (average), 0,876 (minimum), and 1.125(maximum) (Fig. 1F). A total of 262 B-cell epitopes were selected based on the combination of the results (Supplementary Table 3a). BcePred [18] was used to predict B-cell epitopes using accessibility, antigenic propensity, exposed surface, flexibility, hydrophilicity, polarity and turns. Overall, we obtained totally 129 B-cell epitopes (Supplementary Table 3b).

Antigenicity of predicted B-cell epitopes were further evaluated by VaxiJen v2.0 with high stringent threshold of 0.9. Based on transmembrane topology of S protein predicted by TMHMM v2.0, intracellular epitopes were further eliminated. A total of 14 residue length B-cell epitopes were obtained including ‘VRQIAPGQTGKIAD’, ‘VLGQSKRVDFCGKG’, ‘GLTGTGVLTESNKK’ and ‘KIADYNYKLPDDFT’ (Table 1). The four B-cell epitopes were mapped to the 3D structure of SARS-CoV-2 S protein (PDB ID: 6VSB), showing that ‘VLGQSKRVDFCGKG’ and ‘GLTGTGVLTESNKK’ are in less-exposed region (Figure 2 A, and B), while ‘VRQIAPGQTGKIAD’ and ‘KIADYNYKLPDDFT’ locate in the spike head which is the most exposed region (Figure 2 C).

**Table 1.**
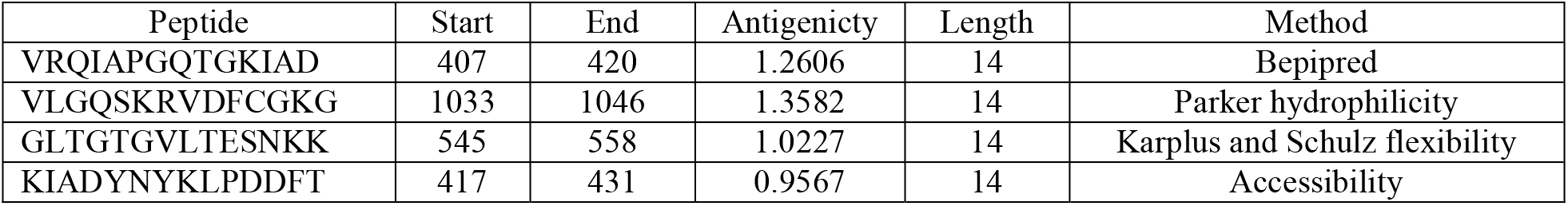
B-cell epitopes present on surface predicted through IEDB and BCPRED along with antigenicity scores.

**Figure 2.**
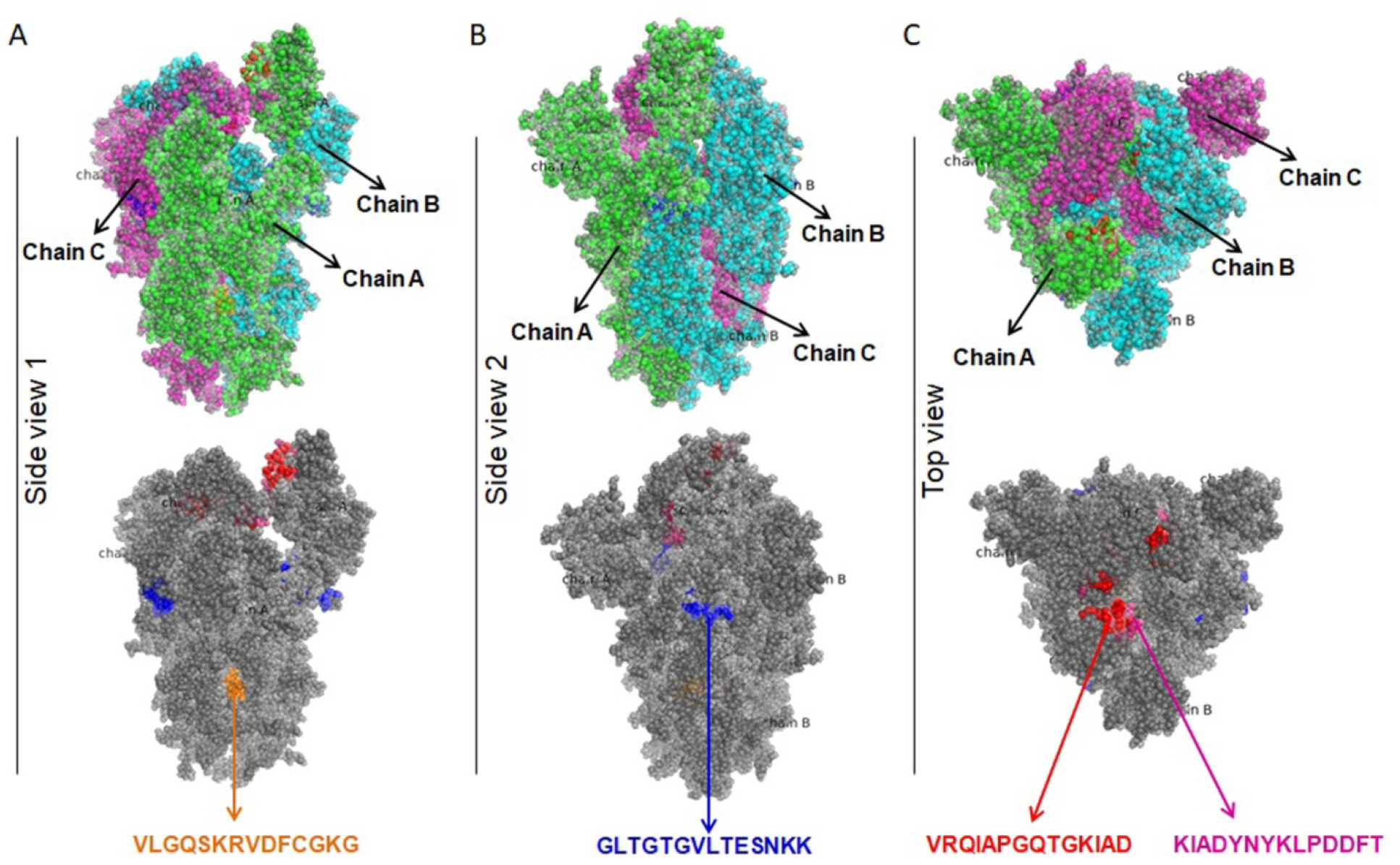
Location of predicited B-cell epitopes on the SARS-CoV-2 S protein (PDB ID 6VSB). A. The location of ‘VLGQSKRVDFCGKG’; B. The location of ‘GLTGTGVLTESNKK’; C. The location of ‘VRQIAPGQTGKIAD’ and ‘KIADYNYKLPDDFT’. Chain A, B, and C are shown in green, cyan, and pink color, respectively.

Discontinuous B-cell epitopes were predicted by Discotope 2.0 using A, B, and C chain of 3D structure of S protein (PDB ID: 6VSB), respectively. The positions of discontinuous epitopes were mapped on the surface of 3D structure of S protein (Figure 3A). Most discontinuous B-cell epitopes were mapped on the fully-exposed ‘spike head’ region (Figure 3B) and less-exposed ‘spike stem’, while a few located in the ‘spike root’ region of the spike (Supplementary Table 3c).

**Figure 3.**
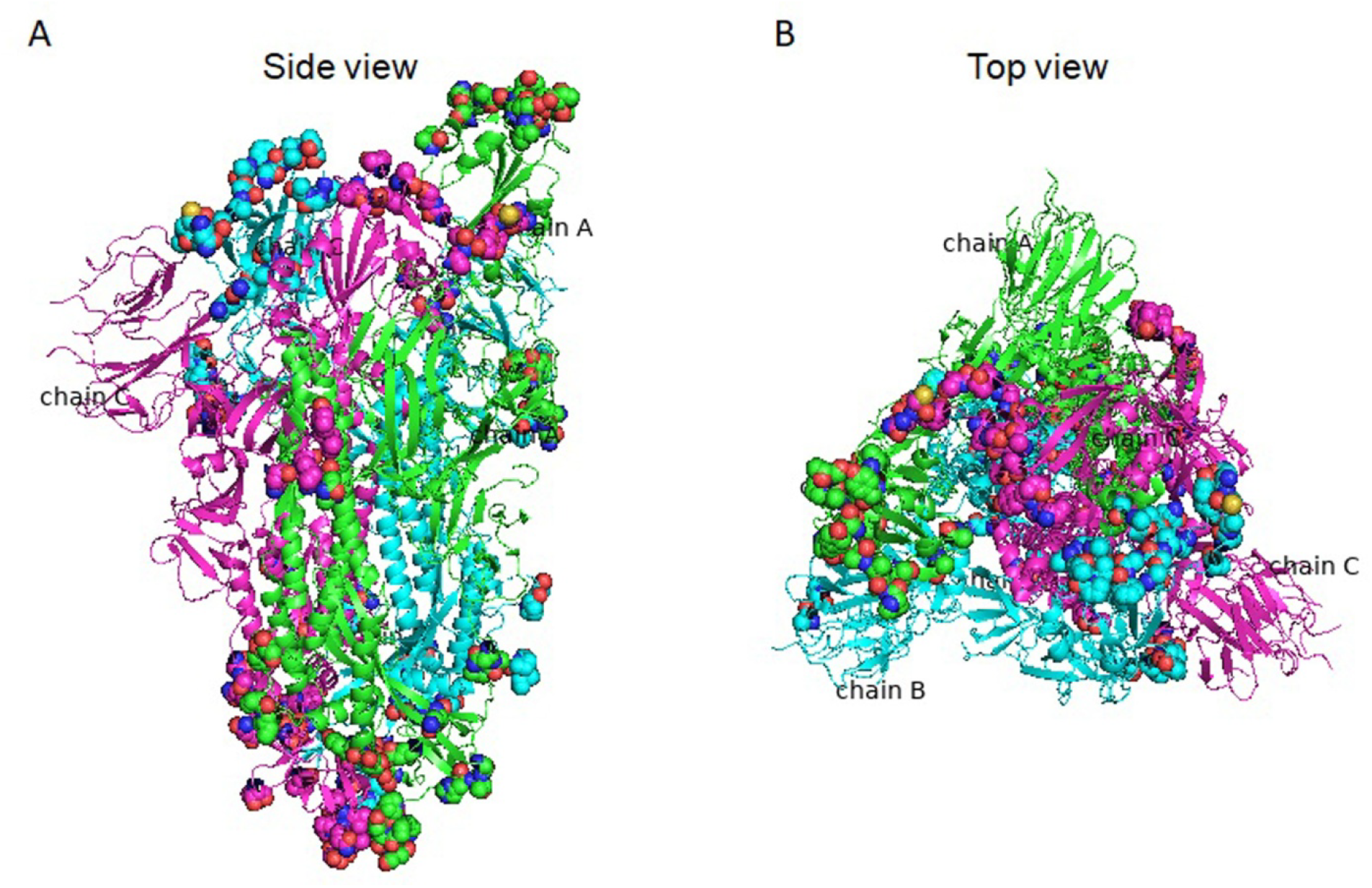
Location of discontinuous B-cell epitopes predicted throough DiscoTope 2 on the 3D structure of the SARS-CoV-2 S protein (PDB ID 6VSB). A. Side-view. B. top-view.

### Allergenicity, toxicity and stability analysis of B-cell epitopes

Allergenicity of B-cell epitopes were assessed by Allergen FP 1.0, leading to the result that all of the four B-cell epitopes were predicted to be not allergenic (Table 2a). Toxicity, hydrophobicity, hydropathicity, hydrophilicity and charge of B-cell epitopes were examined by a support vector machine (SVM) based method, ToxinPred. The result demonstrates that all of B-cell epitopes were predicted to be non-toxin (Table 2a). The stability of B-cell epitopes were evaluated through the number of peptide digesting enzymes by protein digest server. All B-cell epitopes were found to have multiple non-digesting enzymes varying from 4 to 7 enzymes (Table 2b).

**Table 2a.** Allergenicity, toxicity, hydro and physiochemical properties of B-cell epitopes.

| Peptide | Allergenicity | Toxicity | Hydrophobicity | Hydropathicity | Hydrophilicity | Charge | pI | Mol wt |
| --- | --- | --- | --- | --- | --- | --- | --- | --- |
| VRQIAPGQTGKIAD | NA | NT | -0.17 | -0.37 | 0.21 | 1 | 9.1 | 1453.6 |
| VLGQSKRVDFCGKG | NA | NT | -0.21 | -0.27 | 0.3 | 2 | 9.36 | 1494 |
| GLTGTGVLTESNKK | NA | NT | -0.16 | -0.51 | 0.23 | 1 | 8.94 | 1404.8 |
| KIADYNYKLPDDFT | NA | NT | -0.22 | -0.99 | 0.26 | -1 | 4.43 | 1703.1 |

**Table 2b.**
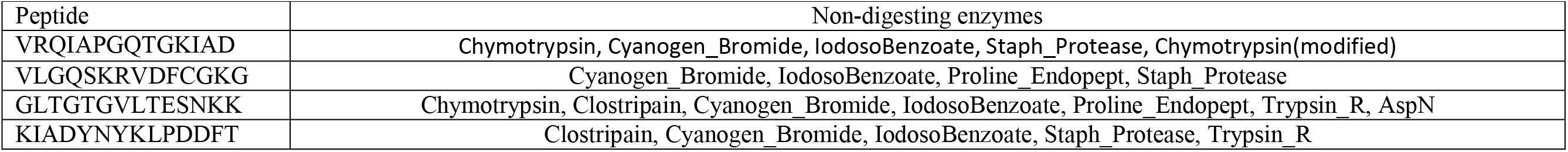
Non-digesting enzymes of B-cell epitopes.

### Identification of T-cell epitopes

Peptide_binding_to_MHC_class_I_molecules tool of IEDB and HLA class I set [20] was utilized to predict T-cell epitopes for S protein. Percentile rank with threshold of 1% was used to filter out peptide-allele with weak binding affinity. The antigenicity score of each peptide was calculated by VaxiJen v2.0 to evaluate its antigenicity. A peptide having both high antigenicity score and capacity to bind with larger number of alleles is considered to have high potentials to initiate a strong defense response. High stringent criteria were used to filter peptides with antigenicity score larger than or equal to 1 and the number of binding alleles larger than or equal to 3. Utilizing the evaluating method above, we obtained a total of 9 MHC class-I allele binding peptides (Table 3). The peptide ‘IPFAMQMAYR’ has the highest antigenicity score of 1.5145 and binds with three alleles including A*68:01, B*35:01, and A*33:01. The peptide ‘FAMQMAYRF’ have the highest number of binding MHC class-I alleles (6 alleles) including B*35:01, B*53:01, A*23:01, B*58:01, A*24:02, and B*08:01 with strong antigenicity score of 1. 0278.

**Table 3.**
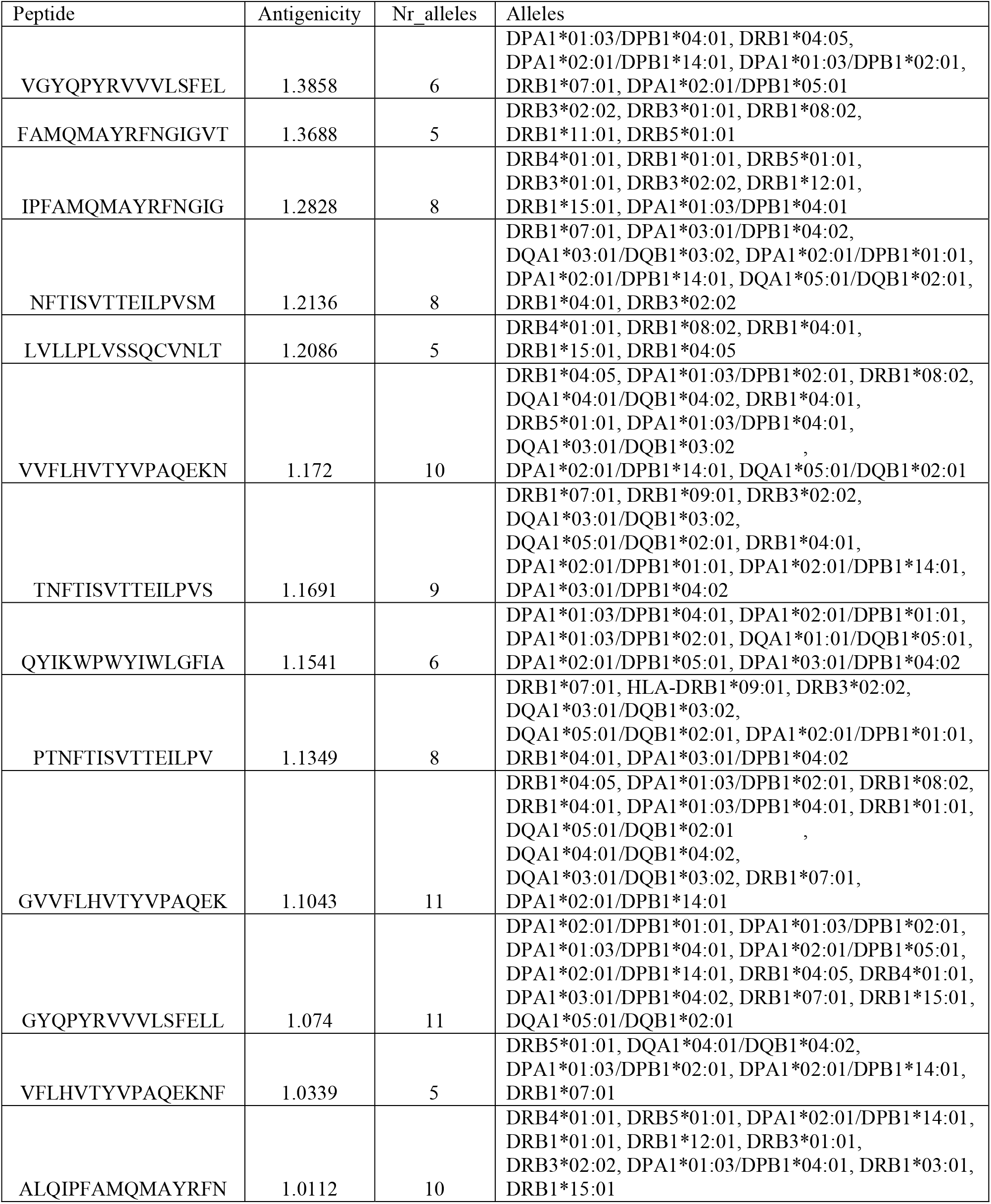
MHC class-I allele binding epitopes.

Peptide_binding_to_MHC_class_II_molecules tool of IEDB and HLA class II set [21] was utilized to predict T-cell epitopes for S protein. Percentile rank with threshold of 10% was used to filter out peptide-allele with weak binding affinity. The antigenicity score of each peptide was calculated by VaxiJen v2.0 to evaluate its antigenicity. A high stringent standard was used to filter peptides with antigenicity score larger than or equal to 1 and the number of binding alleles larger than or equal to 5. As a result, we obtained a total of 13 MHC class-II allele binding peptides (Table 4). The peptide ‘VGYQPYRVVVLSFEL’ has the highest antigenicity score of 1.3858 and binds with six alleles including DPA1*01:03/DPB1*04:01, DRB1*04:05, DPA1*02:01/DPB1*14:01, DPA1*01:03/DPB1*02:01, DRB1*07:01, and DPA1*02:01/DPB1*05:01. The peptides ‘GVVFLHVTYVPAQEK’ and ‘GYQPYRVVVLSFELL’ have the highest number of binding MHC class-II alleles (11 alleles) including DRB1*04:05, DPA1*01:03/DPB1*02:01, DRB1*08:02, DRB1*04:01, DPA1*01:03/DPB1*04:01, DRB1*01:01, DQA1*05:01/DQB1*02:01, DQA1*04:01/DQB1*04:02, DQA1*03:01/DQB1*03:02, DRB1*07:01, DPA1*02:01/DPB1*14:01 for ‘GVVFLHVTYVPAQEK’ with strong antigenicity score of 1.1043 and DPA1*02:01/DPB1*01:01, DPA1*01:03/DPB1*02:01, DPA1*01:03/DPB1*04:01, DPA1*02:01/DPB1*05:01, DPA1*02:01/DPB1*14:01, DRB1*04:05, DRB4*01:01, DPA1*03:01/DPB1*04:02, DRB1*07:01, DRB1*15:01, DQA1*05:01/DQB1*02:01 for ‘GYQPYRVVVLSFELL’ with high antigenicity score of 1.074.

**Table 4.**
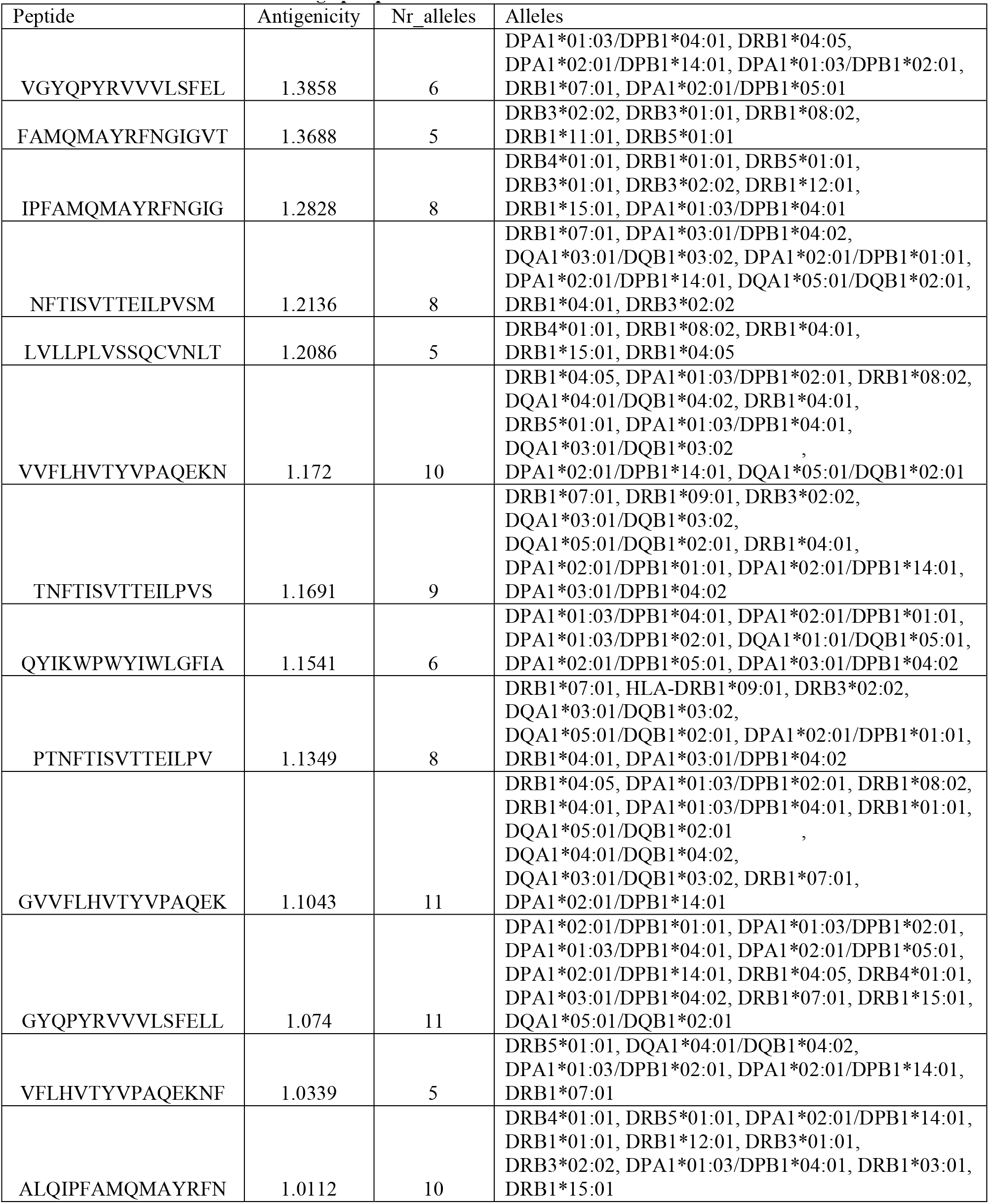
MHC class-II allele binding epitopes.

### Allergenicity, toxicity and stability analysis of T-cell epitopes

Allergenicity of T-cell epitopes were assessed by Allergen FP 1.0. Results showed that two of nine MHC class-I binding peptides were probably non-allergen, and nine of thirteen MHC class-II binding peptides were predicted to be non-allergen (Table 5). Toxicity of T-cell epitopes along with hydrophobicity, hydropathicity, hydrophilicity and charge were evaluated by ToxinPred. All of T-cell epitopes were predicted to be non-toxin (Table 5). The stability of T-cell epitopes were evaluated through the number of peptide digesting enzymes by protein digest server. All T-cell epitopes were found to have multiple non-digesting enzymes varying from 4 to 11 enzymes (Table 6).

**Table 5.**
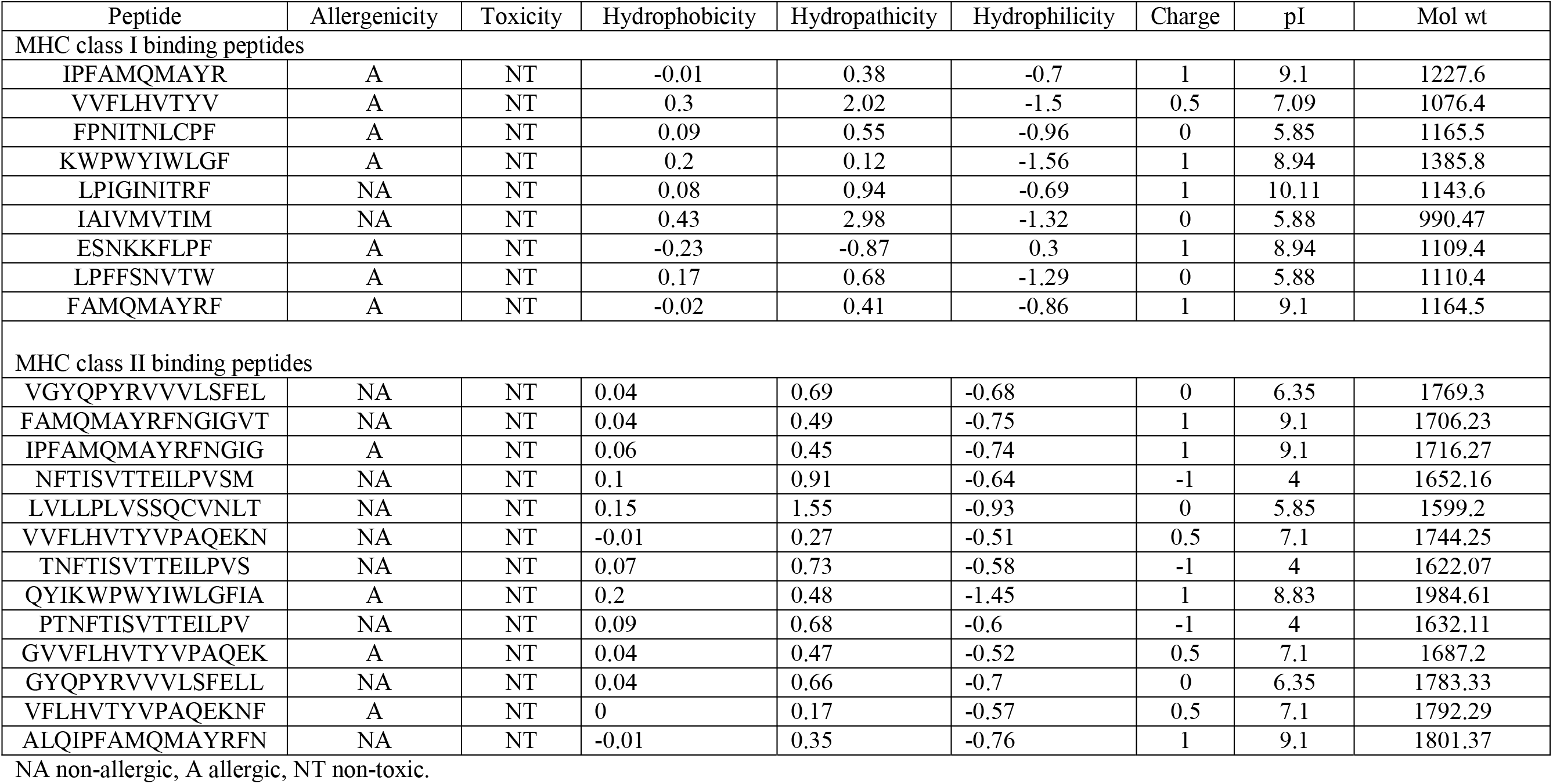
Allergenicity, toxicity, hydro and physiochemical properties of T-cell epitopes.

**Table 6.**
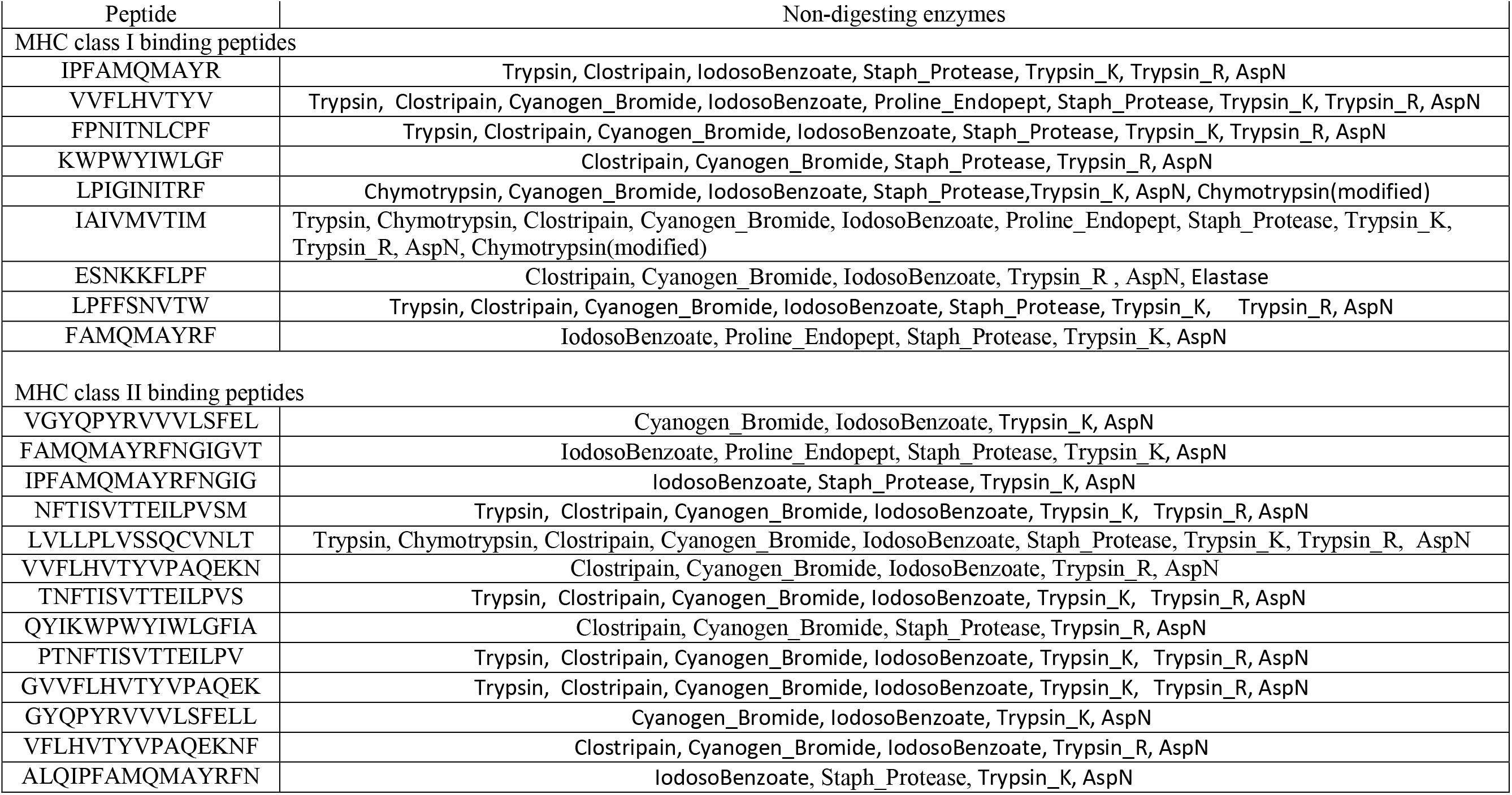
Non-digesting enzymes of T-cell epitopes.

### Interaction of T-cell epitopes with HLA alleles

Protein-peptide interactions are critical in cellular signaling pathways. Two MHC class-I binding epitopes, ‘LPIGINITRF’ and ‘IAIVMVTIM’, were predicted to be non-allergic and non-toxic. Both epitopes were predicted to bind to HLA-B35:01, HLA-B*51:01, and HLA-B*53:01. The 3D structure of human HLA-B35:01(PDB ID: 1A9E) [25], HLA-B*51:01 (PDB ID: 1E27) [26] and HLA-B*53:01 (PDB ID: 1A1O) [27] protein were accessible with co-crystallized peptide in PDB database. Protein-peptide interactions were performed by PepSite [22]. 10 epitope-protein interactions were reported and the top prediction was chosen. HLA-B*35:01 (1A9E) is of a hetero 2mer structure with 386 residues. Epitope ‘LPIGINITRF’ was predicted to significantly bind on the surface of HLA-B35:01(PDB ID: 1A9E) through six hydrogen bonds with Leu-1, Pro-2, Ile-3, Gly-4, Ile-5, and Ans-6 (Figure 4 A). Epitope ‘IAIVMVTIM’ moderately significantly bond to HLA-B35:01(PDB ID: 1A9E) via six hydrogen bonds with Ile-3, Val-4, Met-5, Thr-7, Ile-8, and Met-9 (Figure 4B). Similarly, both epitope ‘LPIGINITRF’ and ‘IAIVMVTIM’ show strong and stable bonding with HLA-B*51:01 (1E27) residues (Figure 4C-D), and HLA-B*53:01 (PDB ID: 1A1O) residues (Figure 4E-F), respectively.

**Figure 4.**
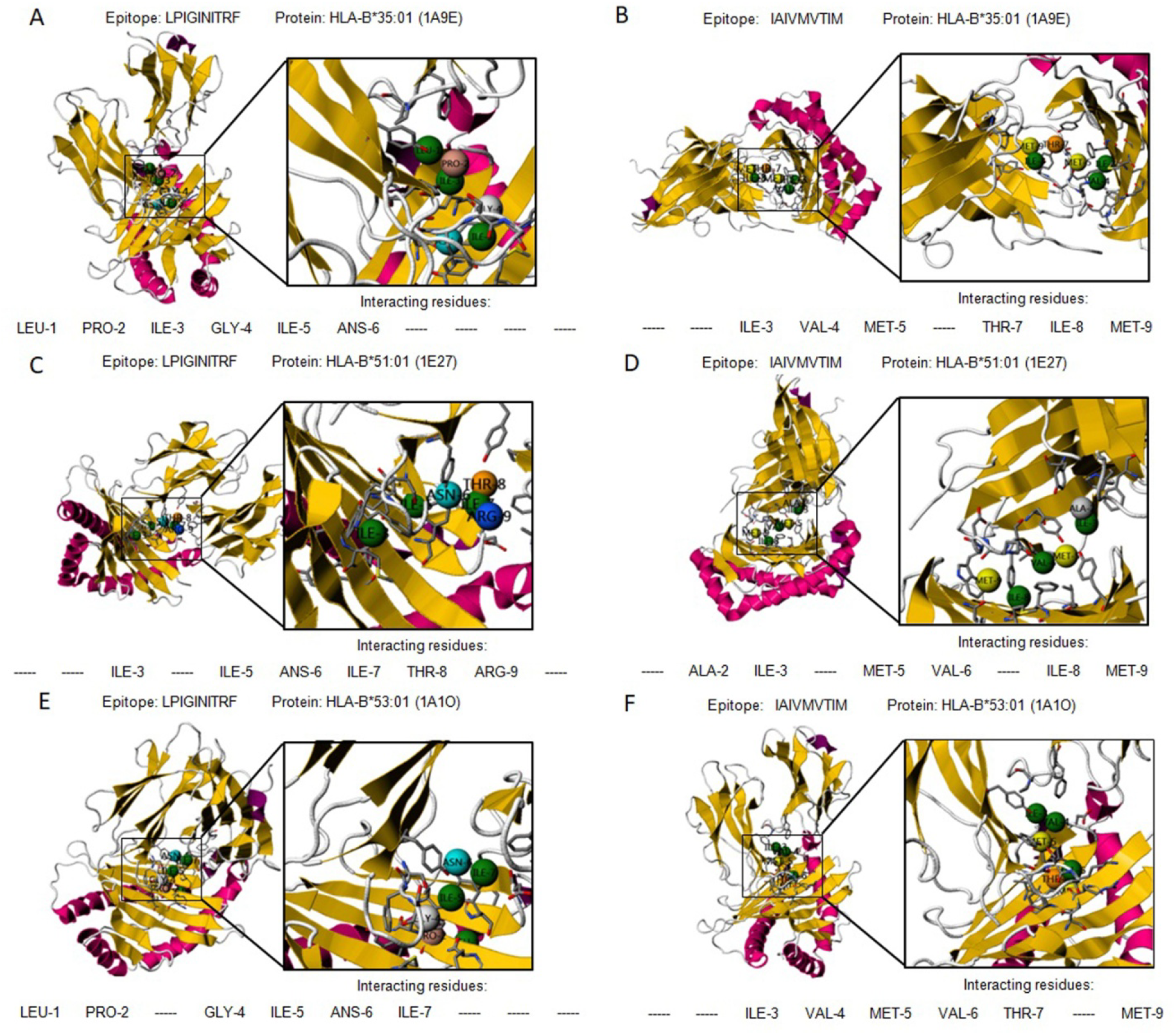
Graphical representation of interactions between Human Leukocyte Antigen and MHC class-I allele binding epitopes. A. LPIGINITRF and HLA-B*35:01 (1A9E); B. IAIVMVTIM and HLA-B*35:01 (1A9E); C. LPIGINITRF and HLA-B*51:01 (1E27); D. IAIVMVTIM and HLA-B*51:01 (1E27);E. LPIGINITRF and HLA-B*53:01 (1A1O); F. IAIVMVTIM and HLA-B*53:01 (1A1O).

### Conservation of B- and T-cell epitopes

Sequence of SARS-CoV-2 S protein with identified mutations from 38 locations worldwide were subjected to multiple sequence alignment of all selected B- and T-cell epitopes. There were no mutations observed occurring in all epitopes, demonstrating all the selected epitopes are conserved in all sequences utilized in the analysis. A phylogenetic tree was generated to show the evolutionary relationship of SARS-CoV-2 S protein collected from 38 locations worldwide (Figure 5).

**Figure 5.**
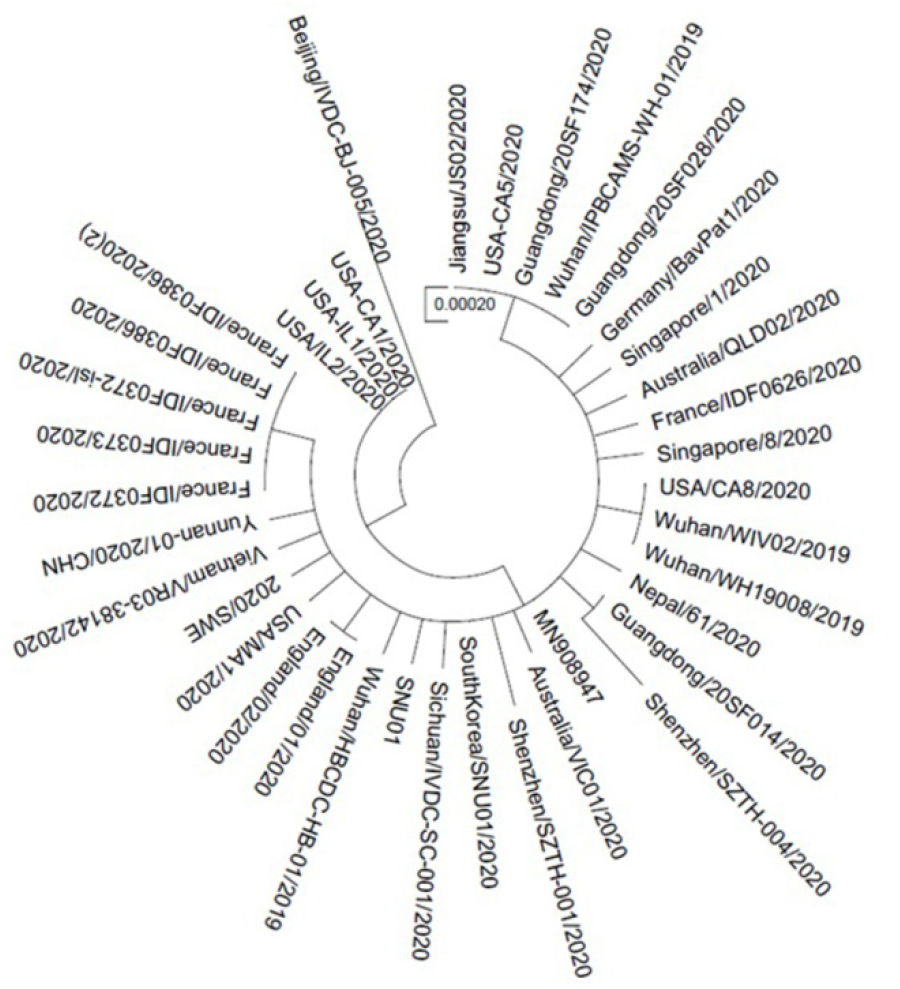
Phylogenetic tree illustrating evolutionary relationships among SARS-CoV-2 isolates from different locations worldwide.

## Discussion

The emergence of SARS-CoV-2 is a serious health threat for the whole society, thus there is an urgent need for drugs and preventative measures. The SARS-CoV-2 infection is characterized by lung infections with symptoms including fever, cough, and shortness of breath. Based on the information from CDC (Centers for Disease Control and Prevention), the symptoms can appear in as few as 2 days or as long as 14 days after exposure to the virus which can transmit from human to human or from contact with infected surfaces and objects [5–7].

It is critical to rapidly identify immune epitopes. The S protein is of crucial in the fuse and entry of virus into host cells [1], therefore it is a primary target for neutralizing antibodies. The specificity of epitope-based vaccines can be enhanced by selecting parts of S protein exposed on the surface [28]. Medical biotechnology is important in developing vaccines against SARS-CoV-2. However, computer-based immune-informatics can improve time and economic effectiveness, as a result, it is also an essential method in immunogenic analysis and vaccine development.

In this study, we characterized the physio-chemical characteristics of SARS-CoV-2 viral genome for epitope candidates and adopted an immune-informatics based pipeline with highly stringent criteria to identify S protein targeted B- and T-cell epitopes that may potentially promote an immune response in the host. The antigenicity, flexibility, solvent accessibility, disulphide bonds of predicted epitopes were evaluated, yielding four potential B-cell epitope and vaccine candidate. Allergenicity and toxicity analysis confirmed the four B-cell epitopes are of non-allergen and non-toxin. Stability analysis revealed that they can not be digested by multiple enzymes. In addition, two MHC class-I and nine MHC class-II binding T-cell epitopes were predicted to interact with numerous HLA alleles and to be highly antigenic in nature. Allergenicity, toxicity, and physiochemical properties of T-cell epitopes were analyzed to increase specificity and selectivity. The stability and safety were confirmed by digestion analysis. All selected B- and T-cell (MHC class-I and II) epitopes were conserved in all isolates of different locations globally without mutations observed yet.

We predict the B- and T-cell epitopes identified here may assist the development of potent peptide-based vaccines to address the SARS-CoV-2 challenge. But the replication of SARS-CoV-2 must be error-prone, which is similar to SARS-CoV with reported mutation rate of 4×10^−4^ substitutions/site/year [29]. Anti-viral vaccines are necessary to be developed before the predicted epitopes are potentially obsolete. In addition, our immune-informatics based pipeline also provides a framework to identify B- and T-cell epitopes having therapeutic potential with excellent scope for SARS-CoV-2, but not limited to specific virus.

## Acknowledgements

This work was supported by grants from the National Natural Science Foundation of China (NSFC No. 11421202, and 11827803 to YBF), the Youth Thousand Scholar Program of China (J.Z.) and Beijing Advanced Innovation Center for Biomedical Engineering, BUAA (J.Z.)

## Authors’ contributions

JZ and YBF conceived and designed this study; JZ, LL, TS, YFH and WDL performed immune-informatics analysis. JZ and YBF wrote the manuscript. JZ, YBF, LL, TS, YFH and WDL improved and revised the manuscript. All authors read and approved the final manuscript.

