## SupplementalFiguresTables for "Epitope-based peptide vaccine design and target site characterization against novel coronavirus disease caused by SARS-CoV-2"

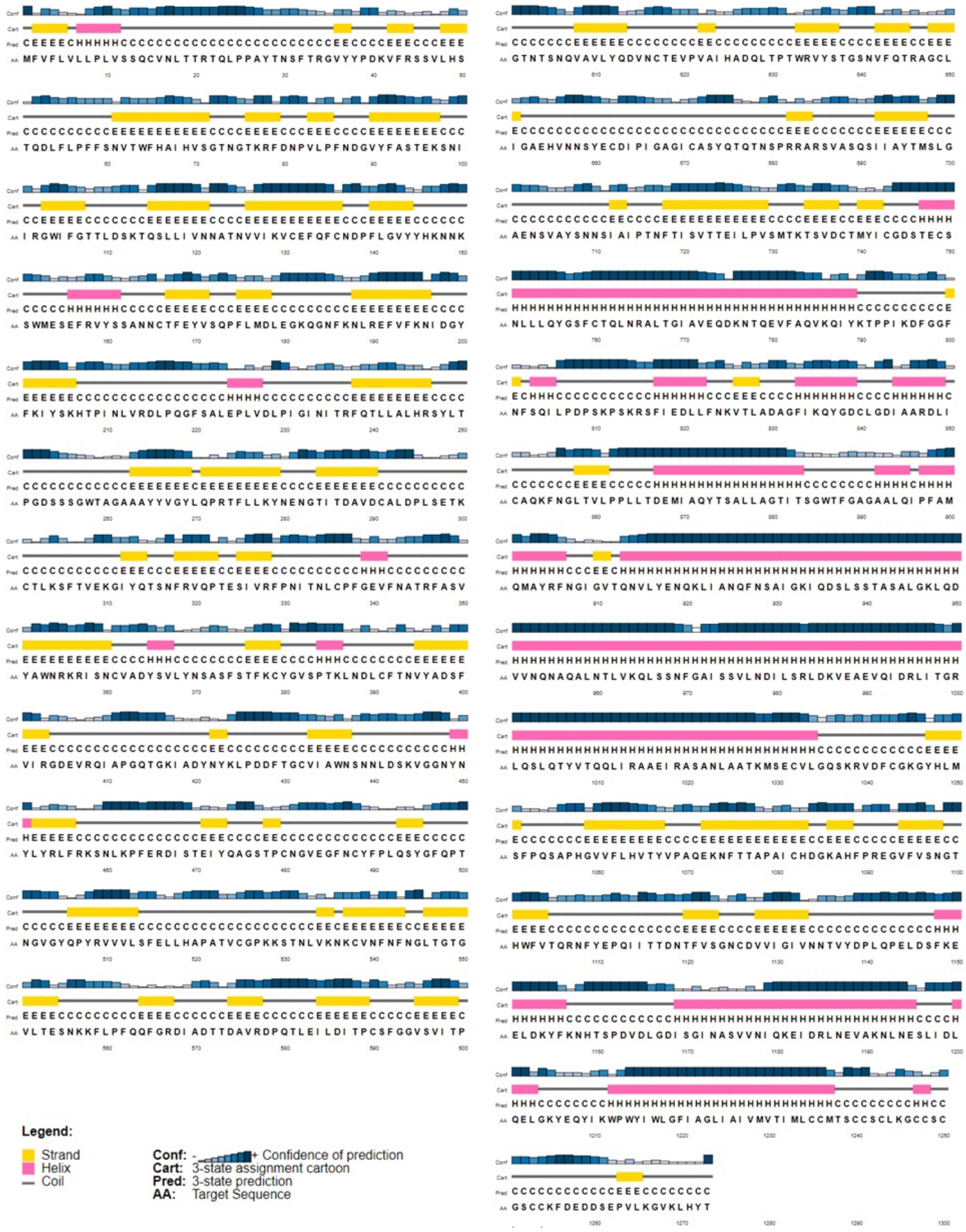


**Supplementary Figure 1**. PSIPRED analysis of the SARS-CoV-2 S protein. Helixes are cylindrical and coloured pink, beta-strands are shown as arrows and coloured yellow, and random coil regions are black.

**Supplementary Table 1:** PhysicoChemical parameters of spike (S) protein computed through ExPASy ProtParam.

| **Parameters** | **SARS-CoV-2 S protein** | | |
|---|---|---|---|
| Mol. Weight | 141178.47 Dalton | | |
| No. of amino acids | 1273 | | |
| Theoretical *pI* | 6.24 | | |
| Instability index (II) | 33.01 (stable) | | |
| No. of Negatively Charged Residues (Asp + Glu) | 110 | | |
| No. of Positively Charged Residues (Arg + Lys) | 103 | | |
| Aliphatic Index | 84.67 | | |
| Grand average of Hydropathicity (GRAVY) | -0.079 | | |
| Atomic Composition | Carbon | 6336 | |
| | Hydrogen | 9770 | |
| | Nitrogen | 1656 | |
| | Oxygen | 1894 | |
| | Sulfur | 54 | |
| Amino Acid Composition | Ala (A) | 79 | 6.2% |
| | Arg (R) | 42 | 3.3% |
| | Asn (N) | 88 | 6.9% |
| | Asp (D) | 62 | 4.9% |
| | Cys (C) | 40 | 3.1% |
| | Gln (Q) | 62 | 4.9% |
| | Glu (E) | 48 | 3.8% |
| | Gly (G) | 82 | 6.4% |
| | His (H) | 17 | 1.3% |
| | Ile (I) | 76 | 6.0% |
| | Leu (L) | 108 | 8.5% |
| | Lys (K) | 61 | 4.8% |
| | Met (M) | 14 | 1.1% |
| | Phe (F) | 77 | 6.0% |
| | Pro (P) | 58 | 4.6% |
| | Ser (S) | 99 | 7.8% |
| | Thr (T) | 97 | 7.6% |
| | Trp (W) | 12 | 0.9% |
| | Tyr (Y) | 54 | 4.2% |
| | Val (V) | 97 | 7.6% |
| | Pyl (O) | 0 | 0% |
| | Sec (U) | 0 | 0% |

**Supplementary Table 2:** Predicted disulphide bonds within residues of S protein by DiANNA 1.1. The weak bonds with lowest Score are highlighted in red.

| Serial no | positions | peptide bonds | scores |
|---|---|---|---|
| 1 | 15 - 1240 | LVSSQCVNLTT – CCMTSCCSCLK | 0.9725 |
| 2 | 131 - 391 | VVIKVCEFQFC – KLNDLCFTNVY | 0.986 |
| 3 | 136 - 662 | CEFQFCNDPFL – NNSYECDIPIG | 0.9966 |
| 4 | 166 - 1236 | SSANNCTFEYV – TIMLCCMTSCC | 0.9983 |
| 5 | 291 - 671 | TDAVDCALDPL – IGAGICASYQT | 0.9094 |
| 6 | 301 - 336 | LSETKCTLKSF – NITNLCPFGEV | 0.9941 |
| 7 | 361 - 488 | KRISNCVADYS – VEGFNCYFPLQ | 0.997 |
| 8 | 379 - 743 | FSTFKCYGVSP – CTMYICGDSTE | 0.9972 |
| 9 | 432 - 1235 | DDFTGCVIAWN – VTIMLCCMTSC | 0.9988 |
| 10 | 480 - 1248 | AGSTPCNGVEG – CLKGCCSCGSC | 0.9987 |
| 11 | 525 - 1247 | APATVCGPKKS – SCLKGCCSCGS | 0.5136 |
| 12 | 538 - 1043 | LVKNKCVNFNF – KRVDFCGKGYH | 0.9995 |
| 13 | 590 - 617 | LDITPCSFGGV – YQDVNCTEVPV | 1 |
| 14 | 649 - 1241 | QTRAGCLIGAE – CMTSCCSCLKG | 0.9997 |
| 15 | 738 - 1243 | KTSVDCTMYIC – TSCCSCLKGCC | 0.0104 |
| 16 | 749 - 1126 | GDSTECSNLLL – FVSGNCDVVIG | 0.9914 |
| 17 | 760 - 1250 | QYGSFCTQLNR – KGCCSCGSCCK | 0.8063 |
| 18 | 840 - 1032 | KQYGDCLGDIA – TKMSECVLGQS | 0.9756 |
| 19 | 851 - 1254 | ARDLICAQKFN – SCGSCCKFDED | 0.9996 |
| 20 | 1082 - 1253 | TAPAICHDGKA – CSCGSCCKFDE | 0.0575 |

**Suppementary Table 3a. B-cell epitopes predicted by IEDB with antigenicity evaluated by VaxiJen.**

| start | end | peptide | vaxijen | method |
|---|---|---|---|---|
| 21 | 31 | RTQLPPAYTNS | 0.871 | Bepipred |
| 71 | 81 | SGTNGTKRFDN | 0.5906 | Bepipred |
| 181 | 186 | GKQGNF | 2.1342 | Bepipred |
| 249 | 261 | LTPGDSSSGWTAG | 0.495 | Bepipred |
| 282 | 287 | NGTITD | 1.1184 | Bepipred |
| 318 | 324 | FRVQPTE | 1.6729 | Bepipred |
| 407 | 420 | VRQIAPGQTGKIAD | 1.2606 | Bepipred |
| 423 | 428 | YKLPDD | -0.9691 | Bepipred |
| 439 | 447 | NNLDSKVGG | 0.8904 | Bepipred |
| 473 | 483 | YQAGSTPCNGV | 0.0881 | Bepipred |
| 495 | 506 | YGFQPTNGVGYQ | 0.7136 | Bepipred |
| 523 | 532 | TVCGPKKSTN | -0.0257 | Bepipred |
| 567 | 580 | RDIADTTDAVRDPQ | 0.44 | Bepipred |
| 597 | 606 | VITPGTNTSN | 0.4217 | Bepipred |
| 675 | 687 | QTQTNSPRRARSV | 0.1763 | Bepipred |
| 772 | 780 | VEQDKNTQE | 0.0684 | Bepipred |
| 788 | 797 | IYKTPPIKDF | -0.1626 | Bepipred |
| 805 | 816 | ILPDPSKPSKRS | 0.5322 | Bepipred |
| 936 | 941 | DSLSST | 0.7417 | Bepipred |
| 1069 | 1077 | PAQEKNFTT | 0.382 | Bepipred |
| 1137 | 1148 | VYDPLQPELDSF | 0.0903 | Bepipred |
| 1157 | 1167 | KNHTSPDVDLG | 1.4039 | Bepipred |
| 1256 | 1265 | FDEDDSEPVL | 0.3154 | Bepipred |
| 13 | 37 | SQCVNLTTRTQLPPAYTNSFTRGVY | 0.686 | Bepipred2.0 |
| 59 | 81 | FSNVTWFHAIHVSGTNGTKRFDN | 0.6767 | Bepipred2.0 |
| 138 | 154 | DPFLGVYYHKNNKSWME | 0.5821 | Bepipred2.0 |
| 177 | 189 | MDLEGKQGNFKNL | 1.2592 | Bepipred2.0 |
| 206 | 221 | KHTPINLVRDLPQGFS | 0.6403 | Bepipred2.0 |
| 250 | 260 | TPGDSSSGWTA | 0.2473 | Bepipred2.0 |
| 304 | 322 | KSFTVEKGIYQTSNFRVQP | 0.5729 | Bepipred2.0 |
| 329 | 363 | FPNITNLCPFGEVFNATRFASVYAWNRKRISNCVA | 0.4466 | Bepipred2.0 |
| 369 | 393 | YNSASFSTFKCYGVSPTKLNDLCFT | 1.4031 | Bepipred2.0 |
| 404 | 426 | GDEVRQIAPGQTGKIADYNYKLP | 1.1017 | Bepipred2.0 |
| 440 | 501 | NLDSKVGGNYNYLYRLFRKSNLKPFERDISTEIYQAGSTPCNGVEGFNCYFPLQSYGFQPTN | 0.3951 | Bepipred2.0 |
| 516 | 536 | ELLHAPATVCGPKKSTNLVKN | 0.0029 | Bepipred2.0 |
| 555 | 562 | SNKKFLPF | 1.3952 | Bepipred2.0 |
| 616 | 632 | NCTEVPVAIHADQLTPT | 0.3987 | Bepipred2.0 |
| 634 | 644 | RVYSTGSNVFQ | -0.1 | Bepipred2.0 |
| 656 | 666 | VNNSYECDIPI | 0.6124 | Bepipred2.0 |
| 672 | 690 | ASYQTQTNSPRRARSVASQ | 0.2556 | Bepipred2.0 |
| 695 | 710 | YTMSLGAENSVAYSNN | 0.6434 | Bepipred2.0 |
| 773 | 779 | EQDKNTQ | 0.1017 | Bepipred2.0 |
| 786 | 800 | KQIYKTPPIKDFGGF | -0.3896 | Bepipred2.0 |
| 807 | 814 | PDPSKPSK | 0.0621 | Bepipred2.0 |
| 828 | 842 | LADAGFIKQYGDCLG | 0.2071 | Bepipred2.0 |
| 1035 | 1043 | GQSKRVDFC | 1.779 | Bepipred2.0 |
| 1107 | 1118 | RNFYEPQIITTD | 0.3529 | Bepipred2.0 |
| 1133 | 1172 | VNNTVYDPLQPELDSFKEELDKYFKNHTSPDVDLGDISGI | 0.1613 | Bepipred2.0 |
| 1252 | 1267 | SCCKFDEDDSEPVLKG | 0.4347 | Bepipred2.0 |
| 4 | 18 | FLVLLPLVSSQCVNL | 0.8302 | Kolaskar and Tongaonkar antigenicity |
| 34 | 41 | RGVYYPDK | 1.0191 | Kolaskar and Tongaonkar antigenicity |
| 44 | 51 | RSSVLHST | 0.5459 | Kolaskar and Tongaonkar antigenicity |
| 53 | 60 | DLFLPFFS | -0.3099 | Kolaskar and Tongaonkar antigenicity |
| 65 | 70 | FHAIHV | 1.6766 | Kolaskar and Tongaonkar antigenicity |
| 81 | 87 | NPVLPFN | 0.5863 | Kolaskar and Tongaonkar antigenicity |
| 115 | 121 | QSLLIVN | 0.8168 | Kolaskar and Tongaonkar antigenicity |
| 125 | 134 | NVVIKVCEFQ | -0.1498 | Kolaskar and Tongaonkar antigenicity |
| 136 | 146 | CNDPFLGVYYH | 0.4109 | Kolaskar and Tongaonkar antigenicity |
| 168 | 174 | FEYVSQP | 0.9073 | Kolaskar and Tongaonkar antigenicity |
| 210 | 216 | INLVRDL | -0.3198 | Kolaskar and Tongaonkar antigenicity |
| 223 | 230 | LEPLVDLP | -0.3271 | Kolaskar and Tongaonkar antigenicity |
| 239 | 248 | QTLLALHRSY | 0.5596 | Kolaskar and Tongaonkar antigenicity |
| 263 | 270 | AAYYVGYL | 0.5218 | Kolaskar and Tongaonkar antigenicity |
| 272 | 278 | PRTFLLK | -1.3917 | Kolaskar and Tongaonkar antigenicity |
| 288 | 295 | AVDCALDP | 0.773 | Kolaskar and Tongaonkar antigenicity |
| 333 | 339 | TNLCPFG | 1.1812 | Kolaskar and Tongaonkar antigenicity |
| 359 | 371 | SNCVADYSVLYNS | -0.1828 | Kolaskar and Tongaonkar antigenicity |
| 376 | 385 | TFKCYGVSPT | 1.5059 | Kolaskar and Tongaonkar antigenicity |
| 430 | 435 | TGCVIA | 0.4716 | Kolaskar and Tongaonkar antigenicity |
| 488 | 495 | CYFPLQSY | 0.9394 | Kolaskar and Tongaonkar antigenicity |
| 505 | 527 | YQPYRVVVLSFELLHAPATVCGP | 0.4697 | Kolaskar and Tongaonkar antigenicity |
| 592 | 599 | FGGVSVIT | 0.7715 | Kolaskar and Tongaonkar antigenicity |
| 607 | 615 | QVAVLYQDV | 0.265 | Kolaskar and Tongaonkar antigenicity |
| 617 | 627 | CTEVPVAIHAD | 0.0499 | Kolaskar and Tongaonkar antigenicity |
| 647 | 653 | AGCLIGA | 0.1743 | Kolaskar and Tongaonkar antigenicity |
| 667 | 674 | GAGICASY | 0.521 | Kolaskar and Tongaonkar antigenicity |
| 687 | 693 | VASQSII | -0.0188 | Kolaskar and Tongaonkar antigenicity |
| 723 | 730 | TTEILPVS | 1.2071 | Kolaskar and Tongaonkar antigenicity |
| 735 | 741 | SVDCTMY | 1.0932 | Kolaskar and Tongaonkar antigenicity |

| | | | | |
|---|---|---|---|---|
| 750 | 763 | SNLLLQYGSFCTQL | 0.7599 | Kolaskar and Tongaonkar antigenicity |
| 781 | 788 | VFAQVKQI | 0.5854 | Kolaskar and Tongaonkar antigenicity |
| 803 | 808 | SQILPD | -0.1542 | Kolaskar and Tongaonkar antigenicity |
| 837 | 843 | YGDCLGD | -0.5555 | Kolaskar and Tongaonkar antigenicity |
| 847 | 853 | RDLICAQ | 1.1443 | Kolaskar and Tongaonkar antigenicity |
| 858 | 864 | LTVLPPL | 0.6786 | Kolaskar and Tongaonkar antigenicity |
| 873 | 880 | YTSALLAG | 0.3798 | Kolaskar and Tongaonkar antigenicity |
| 959 | 966 | LNTLVKQL | -0.7591 | Kolaskar and Tongaonkar antigenicity |
| 973 | 979 | ISSVLND | 0.0414 | Kolaskar and Tongaonkar antigenicity |
| 1002 | 1011 | QSLQTYVTQQ | -0.1009 | Kolaskar and Tongaonkar antigenicity |
| 1030 | 1037 | SECVLGQS | -0.011 | Kolaskar and Tongaonkar antigenicity |
| 1057 | 1070 | PHGVVFLHVTYVPA | 0.8058 | Kolaskar and Tongaonkar antigenicity |
| 1079 | 1085 | PAICHDG | -1.01 | Kolaskar and Tongaonkar antigenicity |
| 1123 | 1132 | SGNCDVVIGI | 0.7421 | Kolaskar and Tongaonkar antigenicity |
| 1174 | 1179 | ASVVNI | 0.8671 | Kolaskar and Tongaonkar antigenicity |
| 1221 | 1256 | IAGLIAIVMVTIMLCCMTSCCSCLKGCCSCGSCCKF | 0.143 | Kolaskar and Tongaonkar antigenicity |
| 1262 | 1270 | EPVLKGVKL | 1.2301 | Kolaskar and Tongaonkar antigenicity |
| 14 | 20 | QCVNLTT | 1.5916 | Parker hydrophilicity |
| 28 | 33 | YTNSFT | -0.7122 | Parker hydrophilicity |
| 47 | 52 | VLHSTQ | 0.8948 | Parker hydrophilicity |
| 70 | 80 | VSGTNGTKRFD | 0.8493 | Parker hydrophilicity |
| 94 | 100 | STEKSNI | 0.6662 | Parker hydrophilicity |
| 109 | 115 | TLDSKTQ | 1.2584 | Parker hydrophilicity |
| 145 | 153 | YHKNNKSWM | 0.3388 | Parker hydrophilicity |
| 161 | 167 | SSANNCT | -0.4197 | Parker hydrophilicity |
| 180 | 188 | EGKQGNFKN | 1.1232 | Parker hydrophilicity |
| 248 | 263 | YLTPGDSSSGWTAGAA | 0.465 | Parker hydrophilicity |
| 280 | 292 | NENGTITDAVDCA | 0.5014 | Parker hydrophilicity |
| 295 | 302 | PLSETKCT | 1.2573 | Parker hydrophilicity |
| 316 | 325 | SNFRVQPTES | 1.2078 | Parker hydrophilicity |
| 356 | 364 | KRISNCVAD | 0.0005 | Parker hydrophilicity |
| 381 | 388 | GVSPTKLN | 1.9197 | Parker hydrophilicity |
| 404 | 409 | GDEVRQ | 0.6701 | Parker hydrophilicity |
| 411 | 425 | APGQTGKIADYNYKL | 1.4441 | Parker hydrophilicity |
| 437 | 449 | NSNNLDSKVGGNY | 0.6657 | Parker hydrophilicity |
| 464 | 470 | FERDIST | -1.2261 | Parker hydrophilicity |
| 472 | 485 | IYQAGSTPCNGVEG | -0.0612 | Parker hydrophilicity |
| 496 | 507 | GFQPTNGVGYQP | 0.6299 | Parker hydrophilicity |
| 522 | 535 | ATVCGPKKSTNLVK | 0.0422 | Parker hydrophilicity |
| 550 | 557 | GVLTESNK | 0.7779 | Parker hydrophilicity |
| 565 | 581 | FGRDIADTTDAVRDPQT | 0.0859 | Parker hydrophilicity |
| 599 | 607 | TPGTNTSNQ | 0.5029 | Parker hydrophilicity |
| 614 | 620 | DVNCTEV | 2.2015 | Parker hydrophilicity |
| 637 | 647 | STGSNVFQTRA | 0.502 | Parker hydrophilicity |
| 654 | 662 | EHVNNSYEC | 1.068 | Parker hydrophilicity |
| 672 | 689 | ASYQTQTNSPRRARSVAS | 0.2963 | Parker hydrophilicity |
| 699 | 712 | LGAENSVAYSNNSI | 0.608 | Parker hydrophilicity |
| 731 | 738 | MTKTSVDC | 1.5932 | Parker hydrophilicity |
| 743 | 750 | CGDSTECS | 0.0977 | Parker hydrophilicity |
| 771 | 781 | AVEQDKNTQEV | 0.3969 | Parker hydrophilicity |
| 806 | 814 | LPDPSKPSK | -0.1058 | Parker hydrophilicity |
| 837 | 842 | YGDCLG | -0.4941 | Parker hydrophilicity |
| 926 | 944 | QFNSAIGKIQDSLSSTASA | 0.3983 | Parker hydrophilicity |
| 949 | 958 | QDVVNQNAQA | 0.2037 | Parker hydrophilicity |
| 966 | 972 | LSSNFGA | 0.6114 | Parker hydrophilicity |
| 1033 | 1046 | VLGQSKRVDFCGKG | 1.3582 | Parker hydrophilicity |
| 1053 | 1058 | PQSAPH | 0.141 | Parker hydrophilicity |
| 1069 | 1077 | PAQEKNFTT | 0.382 | Parker hydrophilicity |
| 1081 | 1091 | ICHDGKAHFPR | -0.8542 | Parker hydrophilicity |
| 1114 | 1128 | IITTDNTFVSGNCDV | 0.1092 | Parker hydrophilicity |
| 1141 | 1154 | LQPELDSFKEELDK | -0.7047 | Parker hydrophilicity |
| 1156 | 1168 | FKNHTSPDVDLGD | 1.0616 | Parker hydrophilicity |
| 1187 | 1194 | NEVAKNLN | -0.0646 | Parker hydrophilicity |
| 1238 | 1264 | TSCCSCLKGCCSCGSCCKFDEDDSEPV | 0.1337 | Parker hydrophilicity |
| 18 | 32 | LTTRTQLPPAYTNSF | 0.79 | Emini surface accessibility |
| 35 | 43 | GVYYPDKVF | 0.0652 | Emini surface accessibility |
| 73 | 80 | TNGTKRFD | 0.2041 | Emini surface accessibility |
| 110 | 115 | LDSKTQ | 1.3071 | Emini surface accessibility |
| 144 | 153 | YYHKNNKSWM | 0.3777 | Emini surface accessibility |
| 179 | 185 | LEGKQGN | 1.8367 | Emini surface accessibility |
| 202 | 208 | KIYSKHT | 0.7773 | Emini surface accessibility |
| 250 | 255 | TPGDSS | 0.3268 | Emini surface accessibility |
| 278 | 284 | KYNENGT | 0.9414 | Emini surface accessibility |
| 314 | 323 | QTSNFRVQPT | 1.405 | Emini surface accessibility |
| 352 | 357 | AWNRKR | 1.632 | Emini surface accessibility |
| 419 | 428 | ADYNYKLPDD | 0.6956 | Emini surface accessibility |
| 437 | 442 | NSNNLD | 1.1859 | Emini surface accessibility |
| 455 | 468 | LFRKSNLKPFERDI | 0.361 | Emini surface accessibility |
| 495 | 500 | YGFQPT | 1.6231 | Emini surface accessibility |
| 569 | 581 | IADTTDAVRDPQT | 0.3018 | Emini surface accessibility |
| 601 | 606 | GTNTSN | 1.2831 | Emini surface accessibility |

| | | | | |
|---|---|---|---|---|
| 627 | 636 | DQLTPTWRVY | 0.6489 | Emini surface accessibility |
| 655 | 660 | HVNNSY | 1.0629 | Emini surface accessibility |
| 674 | 685 | YQTQTNSPRRAR | 0.076 | Emini surface accessibility |
| 773 | 779 | EQDKNTQ | 0.1017 | Emini surface accessibility |
| 786 | 794 | KQIYKTPPI | 0.2705 | Emini surface accessibility |
| 808 | 817 | DPSKPSKRSF | 0.8148 | Emini surface accessibility |
| 914 | 920 | NVLYENQ | 0.4689 | Emini surface accessibility |
| 1068 | 1076 | VPAQEKNFT | 1.0107 | Emini surface accessibility |
| 1105 | 1111 | TQRNFYE | 0.219 | Emini surface accessibility |
| 1139 | 1162 | DPLQPELDSFKEELDKYFKNHTSP | -0.3378 | Emini surface accessibility |
| 1179 | 1186 | IQKEIDRL | -1.0253 | Emini surface accessibility |
| 1202 | 1210 | ELGKYEQYI | 0.5415 | Emini surface accessibility |
| 1256 | 1261 | FDEDDS | -0.0685 | Emini surface accessibility |
| 23 | 34 | QLPPAYTNSFTR | -0.0689 | Chou and Fasman beta turn |
| 36 | 43 | VYYPDKVF | 0.0301 | Chou and Fasman beta turn |
| 71 | 91 | SGTNGTKRFDNPVLPFNDGVY | 0.3544 | Chou and Fasman beta turn |
| 109 | 115 | TLDSKTQ | 1.2584 | Chou and Fasman beta turn |
| 135 | 142 | FCNDPFLG | 0.1982 | Chou and Fasman beta turn |
| 145 | 152 | YHKNNKSW | 0.4099 | Chou and Fasman beta turn |
| 161 | 167 | SSANNCT | -0.4197 | Chou and Fasman beta turn |
| 181 | 188 | GKQGNFKN | 1.0999 | Chou and Fasman beta turn |
| 248 | 260 | YLTPGDSSSGWTA | 0.627 | Chou and Fasman beta turn |
| 279 | 285 | YNENGTI | 0.508 | Chou and Fasman beta turn |
| 292 | 298 | ALDPLSE | 0.5276 | Chou and Fasman beta turn |
| 313 | 320 | YQTSNFRV | 0.371 | Chou and Fasman beta turn |
| 367 | 374 | VLYNSASF | 0.1765 | Chou and Fasman beta turn |
| 380 | 389 | YGVSPTKLND | 1.4531 | Chou and Fasman beta turn |
| 411 | 430 | APGQTGKIADYNYKLPDDFT | 1.0425 | Chou and Fasman beta turn |
| 436 | 452 | WNSNNLDSKVGGNYNYL | 0.8074 | Chou and Fasman beta turn |
| 474 | 507 | QAGSTPCNGVEGFNCYFPLQSYGFQPTNGVGYQP | 0.5024 | Chou and Fasman beta turn |
| 523 | 535 | TVCGPKKSTNLVK | 0.0426 | Chou and Fasman beta turn |
| 537 | 548 | KCVNFNFNGLTG | 1.6969 | Chou and Fasman beta turn |
| 588 | 594 | TPCSFGG | 1.6632 | Chou and Fasman beta turn |
| 599 | 606 | TPGTNTSN | 0.5249 | Chou and Fasman beta turn |
| 636 | 643 | YSTGSNVF | -0.1992 | Chou and Fasman beta turn |
| 655 | 666 | HVNNSYECDIPI | 0.6752 | Chou and Fasman beta turn |
| 674 | 683 | YQTQTNSPRR | -0.2647 | Chou and Fasman beta turn |
| 705 | 714 | VAYSNNSIAI | 1.0545 | Chou and Fasman beta turn |
| 740 | 750 | MYICGDSTECS | -0.3346 | Chou and Fasman beta turn |
| 789 | 802 | YKTPPIKDFGGFNF | 0.2117 | Chou and Fasman beta turn |
| 804 | 815 | QILPDPSKPSKR | 0.2594 | Chou and Fasman beta turn |
| 835 | 842 | KQYGDCLG | 0.4129 | Chou and Fasman beta turn |
| 926 | 931 | QFNSAI | -0.1194 | Chou and Fasman beta turn |
| 934 | 943 | IQDSLSSTAS | 0.4491 | Chou and Fasman beta turn |
| 966 | 972 | LSSNFGA | 0.6114 | Chou and Fasman beta turn |
| 1033 | 1038 | VLGQSK | 1.2255 | Chou and Fasman beta turn |
| 1040 | 1048 | VDFCGKGYH | 0.375 | Chou and Fasman beta turn |
| 1052 | 1058 | FPQSAPH | -0.3069 | Chou and Fasman beta turn |
| 1082 | 1088 | CHDGKAH | -0.2298 | Chou and Fasman beta turn |
| 1096 | 1101 | VSNGTH | 0.7888 | Chou and Fasman beta turn |
| 1119 | 1128 | NTFVSGNCDV | -0.5646 | Chou and Fasman beta turn |
| 1136 | 1146 | TVYDPLQPELD | 0.3454 | Chou and Fasman beta turn |
| 1155 | 1173 | YFKNHTSPDVDLGDISGIN | 0.88 | Chou and Fasman beta turn |
| 1237 | 1264 | MTSCCSCLKGCCSCGSCCKFDEDDSEPV | 0.1043 | Chou and Fasman beta turn |
| 19 | 34 | TTRTQLPPAYTNSFTR | 0.3477 | Karplus and Schulz flexibility |
| 39 | 47 | PDKVFRSSV | -0.7314 | Karplus and Schulz flexibility |
| 49 | 54 | HSTQDL | 0.4956 | Karplus and Schulz flexibility |
| 71 | 83 | SGTNGTKRFDNPV | 0.3648 | Karplus and Schulz flexibility |
| 94 | 99 | STEKSN | 1.0006 | Karplus and Schulz flexibility |
| 108 | 116 | TTLDSKTQS | 1.0106 | Karplus and Schulz flexibility |
| 147 | 152 | KNNKSW | 0.4346 | Karplus and Schulz flexibility |
| 180 | 190 | EGKQGNFKNLR | 1.0042 | Karplus and Schulz flexibility |
| 205 | 210 | SKHTPI | 0.7317 | Karplus and Schulz flexibility |
| 214 | 220 | RDLPQGF | 1.1398 | Karplus and Schulz flexibility |
| 248 | 258 | YLTPGDSSSGW | 0.624 | Karplus and Schulz flexibility |
| 279 | 287 | YNENGTITD | 0.765 | Karplus and Schulz flexibility |
| 294 | 305 | DPLSETKCTLKS | 0.9204 | Karplus and Schulz flexibility |
| 383 | 389 | SPTKLND | 0.839 | Karplus and Schulz flexibility |
| 403 | 408 | RGDEVR | -2.2527 | Karplus and Schulz flexibility |
| 411 | 417 | APGQTGK | 1.7614 | Karplus and Schulz flexibility |
| 424 | 430 | KLPDDFT | -0.051 | Karplus and Schulz flexibility |
| 437 | 448 | NSNNLDSKVGGN | 0.6962 | Karplus and Schulz flexibility |
| 457 | 471 | RKSNLKPFERDISTE | 0.4847 | Karplus and Schulz flexibility |
| 475 | 484 | AGSTPCNGVE | 0.0244 | Karplus and Schulz flexibility |
| 498 | 503 | QPTNGV | 0.0716 | Karplus and Schulz flexibility |
| 526 | 537 | GPKKSTNLVKNK | 0.4692 | Karplus and Schulz flexibility |
| 545 | 558 | GLTGTGVLTESNKK | 1.0227 | Karplus and Schulz flexibility |
| 564 | 582 | QFGRDIADTTDAVRDPQTL | 0.1344 | Karplus and Schulz flexibility |
| 599 | 607 | TPGTNTSNQ | 0.5029 | Karplus and Schulz flexibility |
| 614 | 619 | DVNCTE | 2.375 | Karplus and Schulz flexibility |
| 627 | 632 | DQLTPT | 0.7329 | Karplus and Schulz flexibility |

| | | | | |
|---|---|---|---|---|
| 637 | 642 | STGSNV | 0.3608 | Karplus and Schulz flexibility |
| 675 | 691 | QTQTNSPRRARSVASQS | 0.1698 | Karplus and Schulz flexibility |
| 744 | 750 | GDSTECS | 0.111 | Karplus and Schulz flexibility |
| 773 | 780 | EQDKNTQE | 0.0364 | Karplus and Schulz flexibility |
| 789 | 798 | YKTPPIKDFG | 0.02 | Karplus and Schulz flexibility |
| 806 | 816 | LPDPSKPSKRS | 0.5972 | Karplus and Schulz flexibility |
| 836 | 841 | QYGDCL | 0.1281 | Karplus and Schulz flexibility |
| 880 | 885 | GTITSG | -0.0576 | Karplus and Schulz flexibility |
| 932 | 943 | GKIQDSLSSTAS | 0.5659 | Karplus and Schulz flexibility |
| 946 | 956 | GKLQDVVNQNA | 0.3966 | Karplus and Schulz flexibility |
| 961 | 969 | TLVKQLSSN | -0.5178 | Karplus and Schulz flexibility |
| 982 | 988 | SRLDKVE | 0.0966 | Karplus and Schulz flexibility |
| 994 | 1011 | DRLITGRLQSLQTYVTQQ | -0.2656 | Karplus and Schulz flexibility |
| 1034 | 1039 | LGQSKR | 2.0011 | Karplus and Schulz flexibility |
| 1070 | 1078 | AQEKNFTTA | 0.7576 | Karplus and Schulz flexibility |
| 1116 | 1125 | TTDNTFVSGN | 0.3035 | Karplus and Schulz flexibility |
| 1134 | 1172 | NNTVYDPLQPELDSFKEELDKYFKNHTSPDVDLGDISGI | 0.1406 | Karplus and Schulz flexibility |
| 1179 | 1187 | IQKEIDRLN | -0.4173 | Karplus and Schulz flexibility |
| 1191 | 1196 | KNLNES | 0.863 | Karplus and Schulz flexibility |
| 1202 | 1207 | ELGKYE | 0.4595 | Karplus and Schulz flexibility |
| 1256 | 1269 | FDEDDSEPVLKGVK | 0.6734 | Karplus and Schulz flexibility |

**Suppementary Table 3b. B-cell epitopes predicted by BCEPRED with antigenicity evaluated by VaxiJen.**

| start | end | peptide | vaxijen | method |
|---|---|---|---|---|
| 17 | 47 | NLTTRTQLPPAYTNSFTRGVYYPDKVFRSS | 0.3475 | Accessibility |
| 71 | 84 | SGTNGTKRFDNPV | 0.3648 | Accessibility |
| 91 | 103 | YFASTEKSNIIR | -0.0942 | Accessibility |
| 108 | 118 | TTLDSKTQSL | 0.99 | Accessibility |
| 142 | 161 | GVYYHKNNKSWMESEFRVY | 0.4302 | Accessibility |
| 177 | 193 | MDLEGKQGNFKNLREF | 0.8263 | Accessibility |
| 200 | 212 | YFKIYSKHTPIN | 0.7893 | Accessibility |
| 268 | 285 | GYLQPRTFLLKYNENGT | 0.4239 | Accessibility |
| 294 | 326 | DPLSETKCTLKSFTVEKGIYQTSNFRVQPTES | 0.6225 | Accessibility |
| 350 | 362 | VYAWNRKRISNC | 0.3253 | Accessibility |
| 383 | 391 | SPTKLNDL | 1.0358 | Accessibility |
| 417 | 431 | KIADYNYKLPDDFT | 0.9567 | Accessibility |
| 437 | 446 | NSNNLDSKV | 0.6019 | Accessibility |
| 447 | 472 | GNYNYLYRLFRKSNLKPFERDISTE | 0.1595 | Accessibility |
| 503 | 512 | VGYQPYRVV | 1.4383 | Accessibility |
| 525 | 538 | CGPKKSTNLVKNK | 0.1411 | Accessibility |
| 552 | 562 | LTESNKKFLP | 0.6681 | Accessibility |
| 573 | 584 | TDAVRDPQTLE | 0.2349 | Accessibility |
| 599 | 609 | TPGTNTSNQV | 0.4224 | Accessibility |
| 654 | 664 | EHVNNSYECD | 0.827 | Accessibility |
| 672 | 689 | ASYQTQTNSPRRARSVA | 0.3628 | Accessibility |
| 771 | 783 | AVEQDKNTQEVF | 0.2192 | Accessibility |
| 806 | 821 | LPDPSKPSKRSFIED | 0.0476 | Accessibility |
| 913 | 924 | QNVLYENQKLI | 0.2774 | Accessibility |
| 982 | 990 | SRLDKVEA | 0.1615 | Accessibility |
| 1000 | 1012 | RLQSLQTYVTQQ | 0.1103 | Accessibility |
| 1035 | 1043 | GQSKRVDF | 1.9298 | Accessibility |
| 1067 | 1080 | YVPAQEKNFTTAP | 0.6578 | Accessibility |
| 1104 | 1114 | VTQRNFYEPQ | 0.6509 | Accessibility |
| 1134 | 1164 | NNTVYDPLQPELDSFKEELDKYFKNHTSPD | -0.2097 | Accessibility |
| 1178 | 1198 | NIQKEIDRLNEVAKNLNESL | -0.0099 | Accessibility |
| 1201 | 1214 | QELGKYEQYIKWP | 0.3709 | Accessibility |
| 1254 | 1265 | CKFDEDDSEPV | 0.2159 | Accessibility |
| 1 | 21 | MFVFLVLLPLVSSQCVNLTT | 0.8317 | Antigenic_Propensity |
| 36 | 63 | VYYPDKVFRSSVLHSTQDLFLPFFSNV | 0.0327 | Antigenic_Propensity |
| 114 | 122 | TQSLLIVN | 0.6517 | Antigenic_Propensity |
| 124 | 148 | TNVVIKVCEFQFCNDPFLGVYYHK | 0.072 | Antigenic_Propensity |
| 166 | 178 | CTFEYVSQPFLM | 0.2955 | Antigenic_Propensity |
| 207 | 219 | HTPINLVRDLPQ | -0.1335 | Antigenic_Propensity |
| 223 | 234 | LEPLVDLPIGI | 0.8222 | Antigenic_Propensity |
| 265 | 274 | YYVGYLQPR | 1.4692 | Antigenic_Propensity |
| 332 | 343 | ITNLCPFGEVF | 0.3094 | Antigenic_Propensity |
| 364 | 372 | DYSVLYNS | 0.1757 | Antigenic_Propensity |
| 374 | 386 | FSTFKCYGVSPT | 0.8097 | Antigenic_Propensity |
| 389 | 397 | DLCFTNVY | 1.8569 | Antigenic_Propensity |
| 449 | 458 | YNYLYRLFR | -0.8692 | Antigenic_Propensity |
| 486 | 496 | FNCYFPLQSY | 0.8219 | Antigenic_Propensity |
| 505 | 520 | YQPYRVVVLSFELLH | 0.9711 | Antigenic_Propensity |
| 584 | 602 | ILDITPCSFGGVSVITPG | 1.1031 | Antigenic_Propensity |
| 610 | 623 | VLYQDVNCTEVPV | 0.619 | Antigenic_Propensity |
| 736 | 745 | VDCTMYICG | -0.7197 | Antigenic_Propensity |
| 748 | 761 | ECSNLLLQYGSFC | 0.8661 | Antigenic_Propensity |
| 857 | 868 | GLTVLPPLLTD | 0.5484 | Antigenic_Propensity |
| 946 | 955 | GKLQDVVNQ | 0.3499 | Antigenic_Propensity |
| 961 | 969 | TLVKQLSS | -0.4822 | Antigenic_Propensity |
| 975 | 983 | SVLNDILS | -0.6172 | Antigenic_Propensity |
| 1001 | 1014 | LQSLQTYVTQQLI | -0.1517 | Antigenic_Propensity |

| | | | | |
|---|---|---|---|---|
| 1028 | 1039 | KMSECVLGQSK | 0.1391 | Antigenic_Propensity |
| 1057 | 1070 | PHGVVFLHVTYVP | 0.9305 | Antigenic_Propensity |
| 1122 | 1135 | VSGNCDVVIGIVN | 0.7251 | Antigenic_Propensity |
| 1136 | 1144 | TVYDPLQP | 0.3135 | Antigenic_Propensity |
| 1227 | 1257 | IVMVTIMLCCMTSCCSCLKGCCSCGSCCKF | 0.0464 | Antigenic_Propensity |
| 1262 | 1274 | EPVLKGVKLHYT | 1.4118 | Antigenic_Propensity |
| 144 | 154 | YYHKNNKSWM | 0.3777 | Exposed_Surface |
| 351 | 361 | YAWNRKRISN | 0.5855 | Exposed_Surface |
| 456 | 464 | FRKSNLKP | 1.1111 | Exposed_Surface |
| 526 | 534 | GPKKSTNL | 0.7072 | Exposed_Surface |
| 552 | 560 | LTESNKKF | 0.7708 | Exposed_Surface |
| 677 | 686 | QTNSPRRAR | -0.118 | Exposed_Surface |
| 772 | 782 | VEQDKNTQEV | 0.2465 | Exposed_Surface |
| 784 | 797 | QVKQIYKTPPIKD | 0.3607 | Exposed_Surface |
| 808 | 818 | DPSKPSKRSF | 0.8148 | Exposed_Surface |
| 1148 | 1161 | FKEELDKYFKNHT | -0.967 | Exposed_Surface |
| 1179 | 1187 | IQKEIDRL | -1.0253 | Exposed_Surface |
| 68 | 77 | IHVSGTNGT | 0.8621 | Flexibility |
| 90 | 98 | VYFASTEK | 0.9206 | Flexibility |
| 108 | 116 | TTLDSKTQ | 1.0912 | Flexibility |
| 247 | 256 | SYLTPGDSS | 0.7324 | Flexibility |
| 351 | 359 | YAWNRKRI | 0.7206 | Flexibility |
| 439 | 447 | NNLDSKVG | 1.2952 | Flexibility |
| 454 | 462 | RLFRKSNL | 0.2128 | Flexibility |
| 522 | 532 | ATVCGPKKST | -0.1151 | Flexibility |
| 550 | 558 | GVLTESNK | 0.7779 | Flexibility |
| 597 | 606 | VITPGTNTS | 0.2753 | Flexibility |
| 672 | 685 | ASYQTQTNSPRRA | 0.107 | Flexibility |
| 742 | 750 | ICGDSTEC | -0.2546 | Flexibility |
| 770 | 779 | IAVEQDKNT | 0.4823 | Flexibility |
| 804 | 816 | QILPDPSKPSKR | 0.2594 | Flexibility |
| 932 | 940 | GKIQDSLS | 0.2746 | Flexibility |
| 1031 | 1040 | ECVLGQSKR | 0.9377 | Flexibility |
| 1254 | 1262 | CKFDEDDS | -0.1581 | Flexibility |
| 70 | 79 | VSGTNGTKR | 0.9855 | Hydrophilicity |
| 250 | 258 | TPGDSSSG | 0.0924 | Hydrophilicity |
| 278 | 288 | KYNENGTITD | 0.9589 | Hydrophilicity |
| 437 | 449 | NSNNLDSKVGGN | 0.6962 | Hydrophilicity |
| 525 | 533 | CGPKKSTN | -0.2855 | Hydrophilicity |
| 568 | 579 | DIADTTDAVRD | 0.8257 | Hydrophilicity |
| 673 | 685 | SYQTQTNSPRRA | 0.0145 | Hydrophilicity |
| 743 | 752 | CGDSTECSN | 0.1786 | Hydrophilicity |
| 771 | 782 | AVEQDKNTQEV | 0.3969 | Hydrophilicity |
| 807 | 817 | PDPSKPSKRS | 0.8374 | Hydrophilicity |
| 1157 | 1166 | KNHTSPDVD | 0.7809 | Hydrophilicity |
| 144 | 161 | YYHKNNKSWMESEFRVY | 0.3439 | Polarity |
| 185 | 197 | NFKNLREFVFKN | 0.1722 | Polarity |
| 201 | 209 | FKIYSKHT | 1.1661 | Polarity |
| 298 | 311 | ETKCTLKSFTVEK | 0.5711 | Polarity |
| 351 | 362 | YAWNRKRISNC | 0.4749 | Polarity |
| 402 | 410 | IRGDEVRQ | -0.6918 | Polarity |
| 452 | 472 | LYRLFRKSNLKPFERDISTE | -0.0464 | Polarity |
| 552 | 561 | LTESNKKFL | 0.3167 | Polarity |
| 679 | 689 | NSPRRARSVA | 0.2502 | Polarity |
| 809 | 821 | PSKPSKRSFIED | 0.3363 | Polarity |
| 982 | 993 | SRLDKVEAEVQ | 0.3878 | Polarity |
| 1013 | 1021 | IRAAEIRA | 0.2791 | Polarity |
| 1082 | 1095 | CHDGKAHFPREGV | -0.4141 | Polarity |
| 1144 | 1161 | ELDSFKEELDKYFKNHT | -0.8623 | Polarity |

| | | | | |
|---|---|---|---|---|
| 1179 | 1192 | IQKEIDRLNEVAK | -0.1773 | Polarity |
| 1201 | 1212 | QELGKYEQYIK | 0.2086 | Polarity |
| 1253 | 1265 | CCKFDEDDSEPV | 0.4603 | Polarity |
| 134 | 142 | QFCNDPFL | -0.243 | Turns |
| 145 | 153 | YHKNNKSW | 0.4099 | Turns |
| 160 | 169 | YSSANNCTF | -0.1036 | Turns |
| 435 | 445 | AWNSNNLDSK | 1.0198 | Turns |
| 538 | 547 | CVNFNFNGL | 1.7985 | Turns |
| 601 | 609 | GTNTSNQV | 0.7179 | Turns |
| 653 | 662 | AEHVNNSYE | 0.8405 | Turns |
| 706 | 714 | AYSNNSIA | 0.5473 | Turns |
| 1156 | 1164 | FKNHTSPD | 0.5003 | Turns |

**Suppementary Table 3c. Discontinuous B-cell epitopes predicted by DiscoTope 2 with threshold of -3.7.**

| chain_id | residue_id | residue_name | contact_number | propensity_score | discotope_score |
|---|---|---|---|---|---|
| A | 281 | GLU | 0 | -3.366 | -2.979 |
| A | 282 | ASN | 7 | -2.664 | -3.162 |
| A | 415 | THR | 0 | -3.819 | -3.38 |
| A | 420 | ASP | 4 | -3.618 | -3.662 |
| A | 449 | TYR | 4 | -0.567 | -0.962 |
| A | 450 | ASN | 11 | -1.78 | -2.841 |
| A | 454 | ARG | 14 | -1.224 | -2.694 |
| A | 491 | PRO | 7 | -0.72 | -1.442 |
| A | 492 | LEU | 15 | -0.95 | -2.565 |
| A | 493 | GLN | 9 | -0.572 | -1.541 |
| A | 494 | SER | 7 | -0.846 | -1.553 |
| A | 496 | GLY | 3 | 0.041 | -0.309 |
| A | 498 | GLN | 4 | 0.68 | 0.142 |
| A | 499 | PRO | 5 | 0.178 | -0.417 |
| A | 500 | THR | 0 | 1.907 | 1.688 |
| A | 503 | VAL | 5 | -1.856 | -2.218 |
| A | 505 | TYR | 8 | -1.528 | -2.272 |
| A | 556 | ASN | 2 | -3.79 | -3.584 |
| A | 558 | LYS | 2 | -1.479 | -1.539 |
| A | 560 | LEU | 2 | -1.137 | -1.236 |
| A | 561 | PRO | 0 | -0.961 | -0.851 |
| A | 562 | PHE | 0 | -2.061 | -1.824 |
| A | 703 | ASN | 4 | -2.02 | -2.248 |
| A | 704 | SER | 3 | -1.469 | -1.645 |
| A | 705 | VAL | 10 | -2.821 | -3.646 |
| A | 793 | PRO | 1 | -2.278 | -2.131 |
| A | 794 | ILE | 1 | -2.5 | -2.327 |
| A | 809 | PRO | 4 | -2.691 | -2.841 |
| A | 810 | SER | 9 | -0.669 | -1.627 |
| A | 914 | ASN | 7 | -1.117 | -1.794 |
| A | 917 | TYR | 9 | -2.702 | -3.426 |
| A | 918 | GLU | 13 | -2.285 | -3.517 |
| A | 1071 | GLN | 9 | -2.775 | -3.491 |
| A | 1099 | GLY | 1 | -3.789 | -3.468 |
| A | 1100 | THR | 0 | -3.877 | -3.431 |
| A | 1101 | HIS | 8 | -2.903 | -3.489 |
| A | 1111 | GLU | 19 | -1.693 | -3.684 |
| A | 1118 | ASP | 4 | -3.016 | -3.129 |
| A | 1140 | PRO | 7 | -0.961 | -1.656 |
| A | 1141 | LEU | 5 | -0.257 | -0.802 |
| A | 1142 | GLN | 7 | 0.318 | -0.523 |
| A | 1143 | PRO | 6 | 1.067 | 0.255 |
| A | 1144 | GLU | 6 | 0.716 | -0.056 |
| A | 1145 | LEU | 5 | 0.162 | -0.431 |
| A | 1146 | ASP | 5 | 0.731 | 0.072 |
| B | 187 | LYS | 9 | -2.398 | -3.158 |
| B | 209 | PRO | 1 | -3.243 | -2.985 |
| B | 281 | GLU | 0 | -4.065 | -3.597 |
| B | 282 | ASN | 6 | -2.337 | -2.758 |
| B | 415 | THR | 0 | -2.448 | -2.167 |

| | | | | | |
|---|---|---|---|---|---|
| B | 417 | LYS | 15 | -2.055 | -3.544 |
| B | 420 | ASP | 5 | -3.2 | -3.407 |
| B | 449 | TYR | 3 | -1.991 | -2.107 |
| B | 460 | ASN | 15 | -0.073 | -1.79 |
| B | 462 | LYS | 2 | -3.079 | -2.955 |
| B | 469 | SER | 1 | -2.505 | -2.332 |
| B | 471 | GLU | 1 | -2.616 | -2.43 |
| B | 472 | ILE | 3 | -3.202 | -3.179 |
| B | 473 | TYR | 11 | -2.078 | -3.104 |
| B | 487 | ASN | 0 | -1.605 | -1.42 |
| B | 488 | CYS | 3 | -2.718 | -2.75 |
| B | 489 | TYR | 6 | -3.009 | -3.353 |
| B | 493 | GLN | 12 | -2.368 | -3.476 |
| B | 494 | SER | 8 | -2.189 | -2.858 |
| B | 496 | GLY | 1 | -0.693 | -0.728 |
| B | 497 | PHE | 18 | -1.153 | -3.091 |
| B | 498 | GLN | 7 | 1.188 | 0.246 |
| B | 499 | PRO | 5 | 0.294 | -0.315 |
| B | 500 | THR | 1 | 2.231 | 1.86 |
| B | 503 | VAL | 6 | -1.621 | -2.124 |
| B | 504 | GLY | 0 | -2.702 | -2.392 |
| B | 505 | TYR | 9 | -1.67 | -2.513 |
| B | 556 | ASN | 2 | -3.692 | -3.497 |
| B | 558 | LYS | 2 | -2.13 | -2.115 |
| B | 560 | LEU | 1 | -3.858 | -3.529 |
| B | 561 | PRO | 0 | -3.986 | -3.528 |
| B | 703 | ASN | 4 | -2.12 | -2.336 |
| B | 704 | SER | 3 | -1.967 | -2.086 |
| B | 705 | VAL | 10 | -2.857 | -3.678 |
| B | 716 | THR | 10 | -2.78 | -3.61 |
| B | 793 | PRO | 0 | -2.542 | -2.25 |
| B | 794 | ILE | 3 | -2.633 | -2.675 |
| B | 809 | PRO | 4 | -3.089 | -3.193 |
| B | 810 | SER | 9 | -1.014 | -1.932 |
| B | 914 | ASN | 6 | -1.369 | -1.901 |
| B | 917 | TYR | 9 | -2.268 | -3.042 |
| B | 918 | GLU | 11 | -2.251 | -3.257 |
| B | 1071 | GLN | 7 | -3.08 | -3.53 |
| B | 1111 | GLU | 19 | -1.343 | -3.373 |
| B | 1114 | ILE | 8 | -2.852 | -3.444 |
| B | 1118 | ASP | 5 | -2.997 | -3.228 |
| B | 1140 | PRO | 8 | -0.677 | -1.519 |
| B | 1141 | LEU | 5 | -0.017 | -0.59 |
| B | 1142 | GLN | 7 | 0.372 | -0.476 |
| B | 1143 | PRO | 6 | 0.629 | -0.134 |
| B | 1144 | GLU | 4 | 0.704 | 0.163 |
| B | 1145 | LEU | 5 | 0.171 | -0.424 |
| B | 1146 | ASP | 4 | 0.724 | 0.181 |
| C | 281 | GLU | 0 | -3.898 | -3.45 |
| C | 282 | ASN | 3 | -2.566 | -2.616 |
| C | 415 | THR | 0 | -3.378 | -2.989 |
| C | 449 | TYR | 3 | -1.844 | -1.977 |

| | | | | | |
|---|---|---|---|---|---|
| C | 460 | ASN | 11 | -0.126 | -1.376 |
| C | 462 | LYS | 1 | -2.996 | -2.767 |
| C | 469 | SER | 5 | -2.411 | -2.708 |
| C | 470 | THR | 12 | -2.535 | -3.624 |
| C | 471 | GLU | 2 | -1.882 | -1.896 |
| C | 487 | ASN | 0 | -1.487 | -1.316 |
| C | 488 | CYS | 0 | -2.512 | -2.223 |
| C | 489 | TYR | 7 | -2.629 | -3.132 |
| C | 493 | GLN | 9 | -2.288 | -3.06 |
| C | 494 | SER | 7 | -2.316 | -2.854 |
| C | 496 | GLY | 1 | -1.754 | -1.668 |
| C | 498 | GLN | 3 | -0.804 | -1.057 |
| C | 503 | VAL | 10 | -2.855 | -3.677 |
| C | 504 | GLY | 0 | -2.423 | -2.145 |
| C | 505 | TYR | 9 | -2.339 | -3.105 |
| C | 556 | ASN | 0 | -3.139 | -2.778 |
| C | 558 | LYS | 1 | -1.819 | -1.725 |
| C | 560 | LEU | 1 | -2.713 | -2.516 |
| C | 561 | PRO | 0 | -3.804 | -3.367 |
| C | 703 | ASN | 3 | -2.135 | -2.235 |
| C | 704 | SER | 3 | -1.544 | -1.711 |
| C | 793 | PRO | 0 | -2.671 | -2.364 |
| C | 794 | ILE | 4 | -2.452 | -2.63 |
| C | 809 | PRO | 4 | -3.118 | -3.22 |
| C | 810 | SER | 9 | -1.295 | -2.181 |
| C | 914 | ASN | 9 | -0.693 | -1.648 |
| C | 917 | TYR | 9 | -2.612 | -3.347 |
| C | 918 | GLU | 12 | -1.782 | -2.957 |
| C | 1071 | GLN | 7 | -2.99 | -3.451 |
| C | 1074 | ASN | 7 | -2.785 | -3.27 |
| C | 1100 | THR | 0 | -3.348 | -2.963 |
| C | 1111 | GLU | 18 | -1.566 | -3.456 |
| C | 1114 | ILE | 8 | -2.926 | -3.51 |
| C | 1118 | ASP | 4 | -3.023 | -3.136 |
| C | 1140 | PRO | 8 | -1.41 | -2.168 |
| C | 1141 | LEU | 3 | -0.304 | -0.614 |
| C | 1142 | GLN | 6 | 0.278 | -0.444 |
| C | 1143 | PRO | 6 | 0.79 | 0.009 |
| C | 1144 | GLU | 4 | 0.433 | -0.077 |
| C | 1145 | LEU | 4 | 0.405 | -0.101 |
| C | 1146 | ASP | 4 | 0.234 | -0.253 |
